# An integrative layer-resolved atlas of the adult human meninges

**DOI:** 10.1101/2024.11.13.623440

**Authors:** Theresa Degenhard, Adam M.R. Groh, Elia Afanasiev, Michael Luo, Moein Yaqubi, Jo Anne Stratton, Kevin Petrecca

## Abstract

The human meninges are a dynamic tri-layered brain border that plays a key role in brain development, CSF homeostasis, immune regulation, and higher-level brain function. The meninges have also been implicated in central nervous system (CNS) pathologies such as infection, autoimmunity, and brain trauma. To understand how the meningeal microenvironment is altered under pathological conditions it is necessary to have a complete understanding of its normotypic cellular architecture and function. To date, there is no complete atlas of the normotypic adult human meninges. By surgically extracting each human meningeal layer during surgery, we generated the first layer-resolved map of all meningeal cell types via an integration of whole cell single cell RNA sequencing, multiplexed error-robust fluorescence *in situ* hybridization (MERFISH), and protein immunolabelling. Since fibroblasts play key roles in meningeal homeostasis yet remain less well-characterised than other meningeal cell types, we deeply phenotyped these cells in all layers. We identified 10 fibroblast subpopulations with unique predicted functions that localise to distinct neuroanatomical niches. Fibroblast interaction analysis in the dura and subarachnoid space (SAS) uncovered novel interactions with vascular cell populations mediated by insulin growth factor signaling. Together, these data serve as a comprehensive resource for future investigations of meningeal function in the healthy and diseased brain.

## Introduction

The meninges are a tri-layered membrane separating the brain from the periphery that plays a key role in brain development, blood-CSF-barrier maintenance, waste clearance, immune regulation, and executive function^1–3^. Although still a nascent area of investigation, meningeal dysfunction is now recognized as an important component in the pathogenesis of many central nervous system (CNS) disorders, including meningitis, chronic traumatic encephalopathy, Alzheimer’s disease, and multiple sclerosis^4–9^. While meningeal immune cells and lymphatics have been the topic of considerable investigation in both healthy and diseased contexts^10,11^, there is comparatively less known about homeostatic and disease-associated phenotypes of meningeal fibroblasts, particularly in humans. Yet, evidence from mice suggests that these unique meningeal cells harbor a variety of specialized functions critical for maintenance of the intra- and inter-layer meningeal microenvironments^12,13^.

The developing mouse meninges contain layer-specific fibroblast subpopulations with unique transcriptomic signatures that can be identified by E14; early differentiation of meningeal fibroblasts into layer-specific subtypes suggests that key functions are enacted by each subpopulation^12^. Distinct meningeal fibroblast subtypes also populate the adult murine meninges^13^. In humans, only one study has profiled fibroblast subpopulations in the leptomeninges using single nuclear RNA sequencing and was not able to determine which subtypes derive from the pia versus the arachnoid^14^. Human dural fibroblasts have also been assessed with single cell RNA sequencing (scRNAseq), but the study did not focus on characterising this cell population^15^. A thorough assessment of human meningeal fibroblast populations is therefore lacking.

Using fresh surgically-resected human dura, arachnoid, and pia, we provide the first complete, layer-resolved atlas of the normotypic adult human meninges. For the first time, we captured and characterised the transcriptomic identity of all cell types within each meningeal layer using whole cell scRNAseq and spatially localised these cell types using MERFISH and protein labelling *in situ*. We then analyzed fibroblasts and discovered a diversity of layer-specific subpopulations with unique functions that reside in specific neuroanatomical niches. Profiling of cell-cell interactions between layer-specific meningeal cell populations uncovered unique Insulin Growth Factor (IGF) signaling-mediated crosstalk between fibroblasts and adjacent endothelial cells in both the dura and SAS. To facilitate data mining of this atlas, we developed an open science website [SingloCell](https://singlocell.openscience.mcgill.ca/) through which the scRNAseq and spatial transcriptomic data can be explored.

## Results

### Human meninges contain diverse fibroblast-like, immune, and vascular cell populations

To study whole cell transcriptomic heterogeneity of the adult human meninges, we surgically extracted six dura mater samples, three arachnoid mater samples, and three pia mater samples from separate patients for immediate scRNAseq (**Figure 1A**). Although tissue samples for scRNAseq and *in situ* tissue validation were sourced far from the tumor site during surgical extraction (**Table 1**; **Supp. Figure 1A**), copy number variation analysis was nevertheless performed to verify that no tumor cells were present in the samples (**Supp. Figure 1B**). Five additional age and sex-matched leptomeningeal and dura samples were collected from separate patients to create a tissue microarray which was used for *in situ* confirmation of scRNAseq findings via MERFISH and protein labelling (**Figure 1A**).

**Figure 1:**
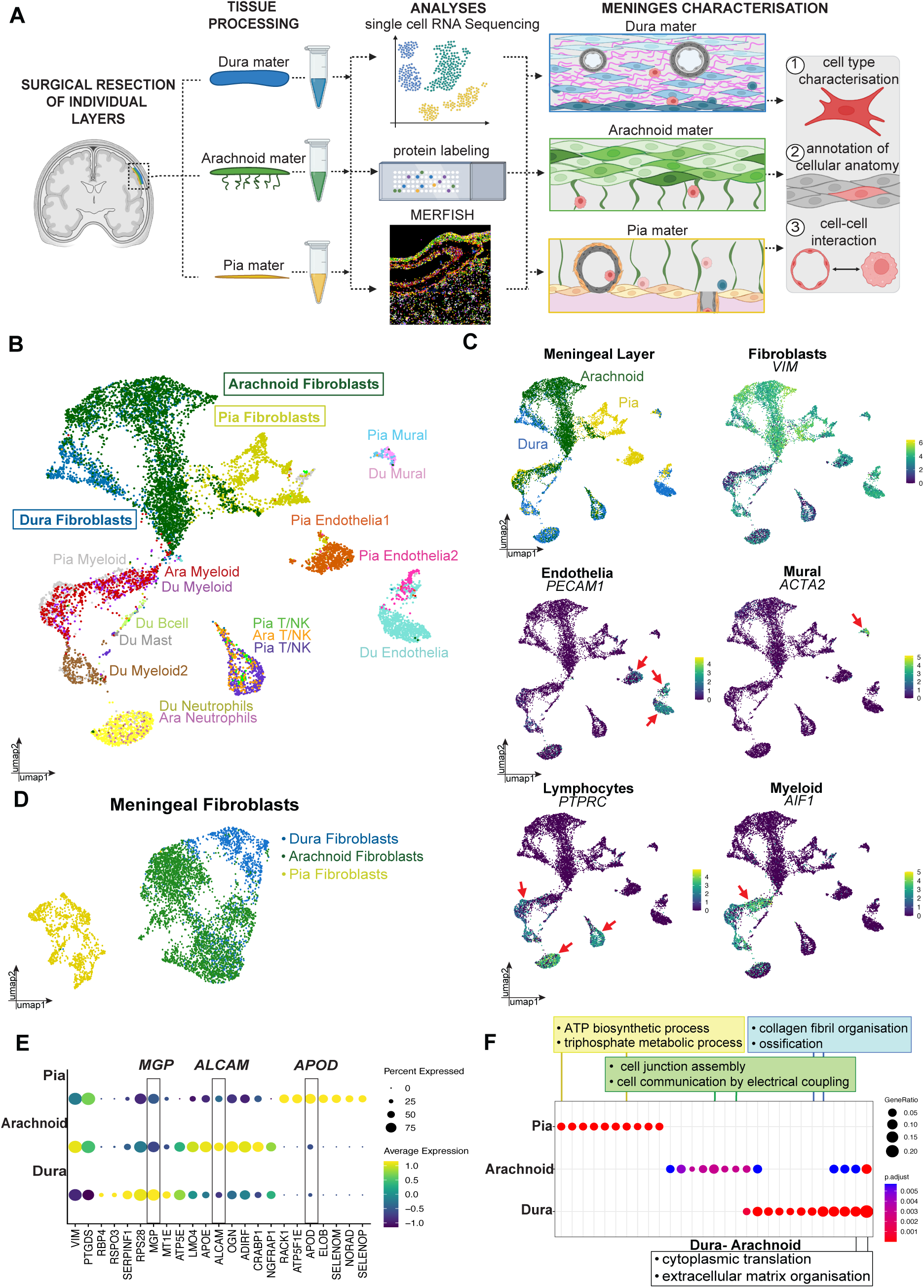
Human meninges contain diverse fibroblast-like, immune, and vascular cell populations. **(A)** Schematic of workflow used to study the cell populations of each meningeal layer. Individual layers (dura, arachnoid and pia) were extracted during brain surgery and immediately processed for single cell RNA sequencing, protein labelling, or MERFISH. These methods allowed for cell type characterisations within each meningeal layer, anatomical localisation of cell types, and predicted cell-cell interactions. **(B)** UMAP of all cell populations captured across all meningeal samples (Table 1). Each cluster is annotated by a unique colour. **(C)** Feature plots demonstrating elevated average expression of marker genes for meningeal fibroblasts (*VIM*-high), myeloid cells (*AIF1*), endothelial cells (*PECAM1*) and mural cells (*ACTA2*). Colour indicates average expression. **(D)** UMAP of meningeal fibroblast populations after subsetting from all meningeal cell types: dura fibroblasts. **(E)** Dotplot demonstrating differential average expression of fibroblast marker genes in each meningeal layer. Dot colour represents average gene expression level and dot size represents the percentage of cells expressing a particular gene. **(F)** GO plot demonstrating the unique and shared biological processes associated with the fibroblasts of each meningeal layer, when compared to one another. Pia fibroblasts showed enrichment of genes associated with proton transmembrane transporter activity, arachnoid fibroblasts showed upregulated genes associated with proteoglycan binding, and dura fibroblasts showed enrichment of genes associated with the extracellular matrix. Arachnoid and pia fibroblasts shared expression of genes associated with cadherin binding, dura and arachnoid fibroblasts highly expressed genes associated with integrin binding, and dura and pia fibroblasts showed shared enrichment of genes related to growth factor binding. Dot colour represents average gene expression level and dot size represents gene ratio.

**Table 1.** Human meninges samples used for single cell RNA sequencing.

| Sample | Tissue | Age | Sex | Pathology | Location |
| --- | --- | --- | --- | --- | --- |
| SD481 | Dura | 45 | f | Meningioma | R, convexity, frontal/parietal lobe |
| SD491 | Dura | 28 | m | Astrocytoma | R, convexity, parietal lobe |
| SD504 | Dura | 55 | f | Ganglion | R, convexity, parietal lobe |
| SD506 | Dura | 66 | f | Meningioma | R, convexity, frontal, parafalcine |
| SD508 | Dura | 49 | f | Glioblastoma | L, convexity, occipital lobe |
| SD509 | Dura | 70 | m | Glioblastoma | R, convexity, temporal lobe |
| SA493 | Arachnoid | 34 | f | Meningioma | R, convexity, posterior, frontal, parafalcine |
| SA506 | Arachnoid | 66 | f | Meningioma | R, convexity, posterior, frontal, parafalcine |
| SA522 | Arachnoid | 59 | f | Meningioma | L, convexity, posterior frontal, parafalcine |
| SP9 | Pia | 66 | m | Glioblastoma | R, convexity, temporal occipital |
| SP10 | Pia | 57 | m | Glioblastoma | R, convexity, temporal occipital |
| SP12 | Pia | 72 | m | Glioblastoma | L, convexity, frontal |

We first profiled the entire cellular landscape of the human meninges by generating a dataset containing all sequenced dura, arachnoid, and pia samples. Nineteen unique cell clusters were identified. These grouped into meningeal or fibroblast-like cells, vascular cells, and immune cells (**Figure 1B**). Fibroblast-like meningeal cells were marked by known high expression of *VIM* and *PTGDS* and a lack of *AIF1, CD31, CD3D and ACTA2* (**Figure 1C; Supp. Figure 2**)^16^. Vascular cells included endothelial cells (*PECAM1*) and mural cells (*ACTA2*) (**Figure 1C**). While mural cells comprised a single cluster, three separate endothelial cell clusters were captured: a dura-derived cluster (Dura endothelial cells; *AQP1*, *PECAM1*); a pia-derived cluster (SAS Endothelia cells1; *CLDN5, PECAM^low^*); and a cluster that included cells from pia and dura (SAS Endothelia cells2; *CLDN5*, *PECAM1*) (**Figure 1B**; **Supp. Figure 2**).

Eleven immune cell clusters were identified: T cells (*CD3D*, *CD8A*); NK cells (*NKG7*, *GZMA*); B cells (*MS4A1*, *CD19*); neutrophils (*S100A9*, *S100A8*); macrophages (*IBA1*, *CSF1R*); monocytes (*CD14*, *FCGR3A*); and mast cells (*CPA3*, *TPSAB1*) (**Figure 1B and C; Supp. Figure 2**). While T/NK cells and myeloid cells were captured in all three meningeal layers, B cells, neutrophils, and mast cells were only found in the arachnoid and/or dura (**Figure 1B; Supp. Figure 2**). Finally, we searched for known markers of lymphatic endothelial cells^17^ but did not find any evidence of this population in our dataset (**Supp. Figure 2**).

While evaluating specific cell proportions using scRNAseq data is subject to considerable technical variation^18^, we have included this data to demonstrate that the majority of cell populations captured in each meningeal layer were present in all samples (**Supp. Figure 3A, B and C**).

Having explored the basic cellular constituents of all three meningeal layers, we wanted to determine whether individual layers of the meninges contained transcriptomically distinct fibroblast-like meningeal cells that might execute different functions depending on their unique neuroanatomical niches. By subsetting out *VIM* and *PTGDS*-positive cells from the integrated meningeal dataset and re-clustering them, we uncovered distinct dura, arachnoid, and pia fibroblast-like cell populations (**Figure 1D**) identified by differential expression of *MGP*, *ALCAM*, and *APOD*, respectively, amongst a variety of additional markers (**Figure 1E**).

By determining the differentially-expressed genes associated with the fibroblast-like cells of each meningeal layer, we predicted layer-specific differences in the function of fibroblast-like cells using gene ontology analysis (**Figure 1F; Supp. Table 1**). This revealed extracellular matrix organization as an over-represented pathway in dura fibroblasts, proteoglycan binding in arachnoid fibroblasts, and proton transmembrane transporter activity in pia fibroblasts (**Figure 1F**). There were also shared over-represented pathways; pia and arachnoid fibroblasts upregulated genes related to cadherin-binding compared to dura fibroblasts. Furthermore, arachnoid and dura fibroblasts upregulated genes related to integrin-binding compared to pia fibroblasts, and pia and dura fibroblasts upregulated genes related to growth factor binding compared to arachnoid fibroblasts (**Figure 1F; Supp. Table 1**).

Lastly, we integrated all meningeal fibroblasts with ependymal cells, another CSF-interfacing barrier cell type, to assess whether this unique neuroanatomical niche conferred any shared transcriptomic identity. We noted transcriptomic similarity between multiciliated ependymal cells and pia fibroblasts. Ependymal cells did not share transcriptomic similarity with arachnoid or dura fibroblasts (**Supp. Figure 4**). This suggests some level of conserved transcriptomic identity that may be related to a shared niche separating the brain from the CSF.

### MERFISH spatial localisation of all primary meningeal cell types in situ

We first assessed the integrity of the meningeal architecture using haematoxylin & eosin staining (**Figure 2A; Supp. Figure 1B**). It showed that the classical features of the meninges were intact: dural periosteal and meningeal layers; dura border cells; arachnoid barrier cells; trabeculae; subarachnoid space (SAS); blood vessel-covering pia; brain-covering pia; and underlying glial limitans superficialis astrocytic end feet (**Figure 2A; Supp. Figure 1B**).

**Figure 2:**
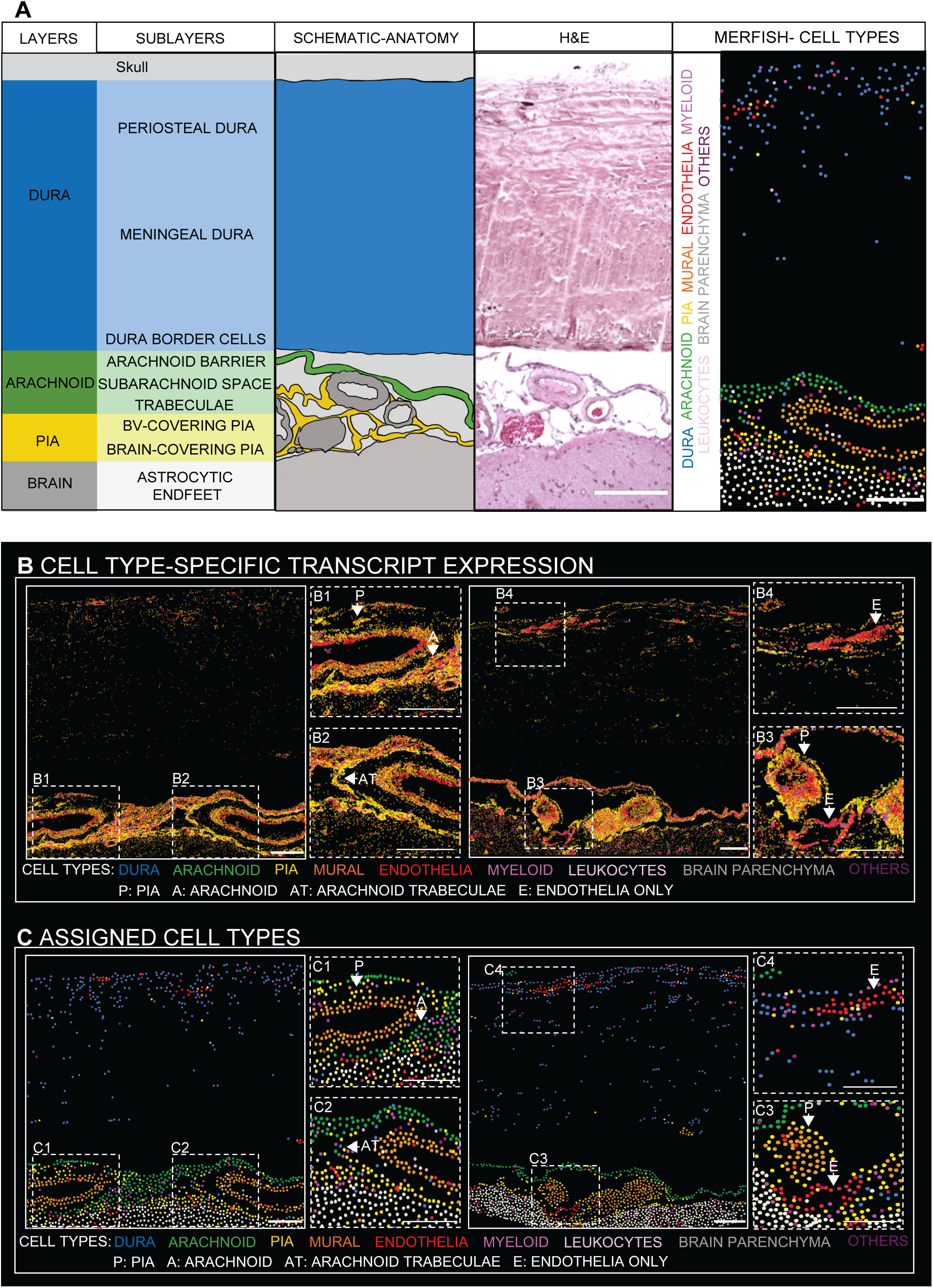
MERFISH confirms the spatial arrangement of meningeal cell types *in situ*. **(A)** Annotated schematic of meningeal anatomy using a representative H&E image and MERFISH panel. Primary neuroanatomical features of the meninges include: periosteal and meningeal layers of the dura; dura border cells; arachnoid barrier cells, subarachnoid space; trabeculae; blood vessel (BV)-covering pia; brain-covering pia; and astrocytic end feet of the glial limitans superficialis. H&E scale bar = 200μm; MERFISH scale bar = 100μm. **(B)** Two representative MERFISH panels from two samples demonstrating the spatial arrangement of primary cell types by visualizing all transcripts associated with cell type signatures. Dotted white boxes B1-4 indicate magnified regions. Each cell type is represented by a different colour. White arrows point to contact points of meningeal fibroblasts and vascular cells. P = Pia; A = Arachnoid; AT = Arachnoid Trabeculae; E = Endothelia only. Scale bars = 100μm. **(C)** Two representative MERFISH panels from two samples demonstrating the spatial arrangement of primary cell types by visualizing cell types following cluster identification and segmentation. Dotted white boxes C1-4 indicate magnified regions. Each cell type is represented by a different colour. White arrows point to contact points of meningeal fibroblasts and vascular cells. P = Pia; A = Arachnoid; AT = Arachnoid Trabeculae; E = Endothelia only. Scale bars = 100μm.

We first used marker genes for layer-specific fibroblasts identified in the scRNAseq analysis to localise these cell populations *in situ* using MERFISH: *MGP* (dura), *ALCAM* (arachnoid), and *APOD* (pia) (**Supp. Figure 5**). We subsequently visualized localisation of transcripts belonging to cell-type specific gene signatures (**Figure 2B**) and localisation of cells derived from segmentation (**Figure 2C**). This allowed visualization of layer-specific fibroblasts amongst other cell types in their microenvironment: mural cells; endothelial cells; myeloid cells; lymphocytes; and underlying glial limitans superficialis astrocytes (**Figure 2B and C**). Protein labelling supported the MERFISH findings (**Supp. Figure 6; Supp. Figure 7**).

Using segmented cells (**Figure 2C**), we then described neuroanatomical relationships between primary meningeal cell subtypes. We primarily noted distinct relationships between meningeal fibroblasts and different sized vessels in the SAS, dura, and brain parenchyma (**Figure 2B**). Larger vessels in the SAS, composed of endothelia and mural cells, were wrapped by blood vessel- and brain-covering pia fibroblasts and arachnoid fibroblasts, in close proximity (**Figure 2, panel C1/2**). We also found examples of adjacent pia and arachnoid fibroblasts that span the SAS, suggesting they form the arachnoid trabeculae (**Figure 2C, panel C2**). Smaller vessels formed by a single endothelia cell layer were only found near the brain surface. These vessels were wrapped by brain-covering pia fibroblasts which sat underneath larger vessels in the SAS (**Figure 2C, panel C3**). In the dura mater, endothelia-lined vessels, largely lacking mural cells, were primarily located in the periosteal layer and were uniquely surrounded by a large quantity of myeloid cells (**Figure 2C, panel C4**). Myeloid cells and some leukocytes were also found throughout the SAS (**Figure 2C**), although they did not appear to congregate perivascularly, as in the dura.

The complete three hundred gene panel used for MERFISH (**Supp. Table 2**) can be interrogated for further evaluation of gene expression across the meningeal layers at the online interactive application [SingloCell](https://singlocell.openscience.mcgill.ca/).

### Characterisation and spatial localisation of dura fibroblast subpopulations

Having spatially resolved the primary layers of the meninges, we profiled intra-layer cellular heterogeneity. Analysis of six dura mater samples revealed nine cell populations: dura fibroblasts; T/NK cells; B cells; neutrophils; myeloid cells (myeloid cells1, myeloid cells2, myeloid cells3); endothelial cells; and mural cells (**Figure 3A and B**). Cell type proportions were estimated by subsampling one thousand cells per sample to avoid bias through sampling differences (**Supp. Figure 3A**). Most cell types were found in all samples (**Supp. Figure 3A**).

**Figure 3:**
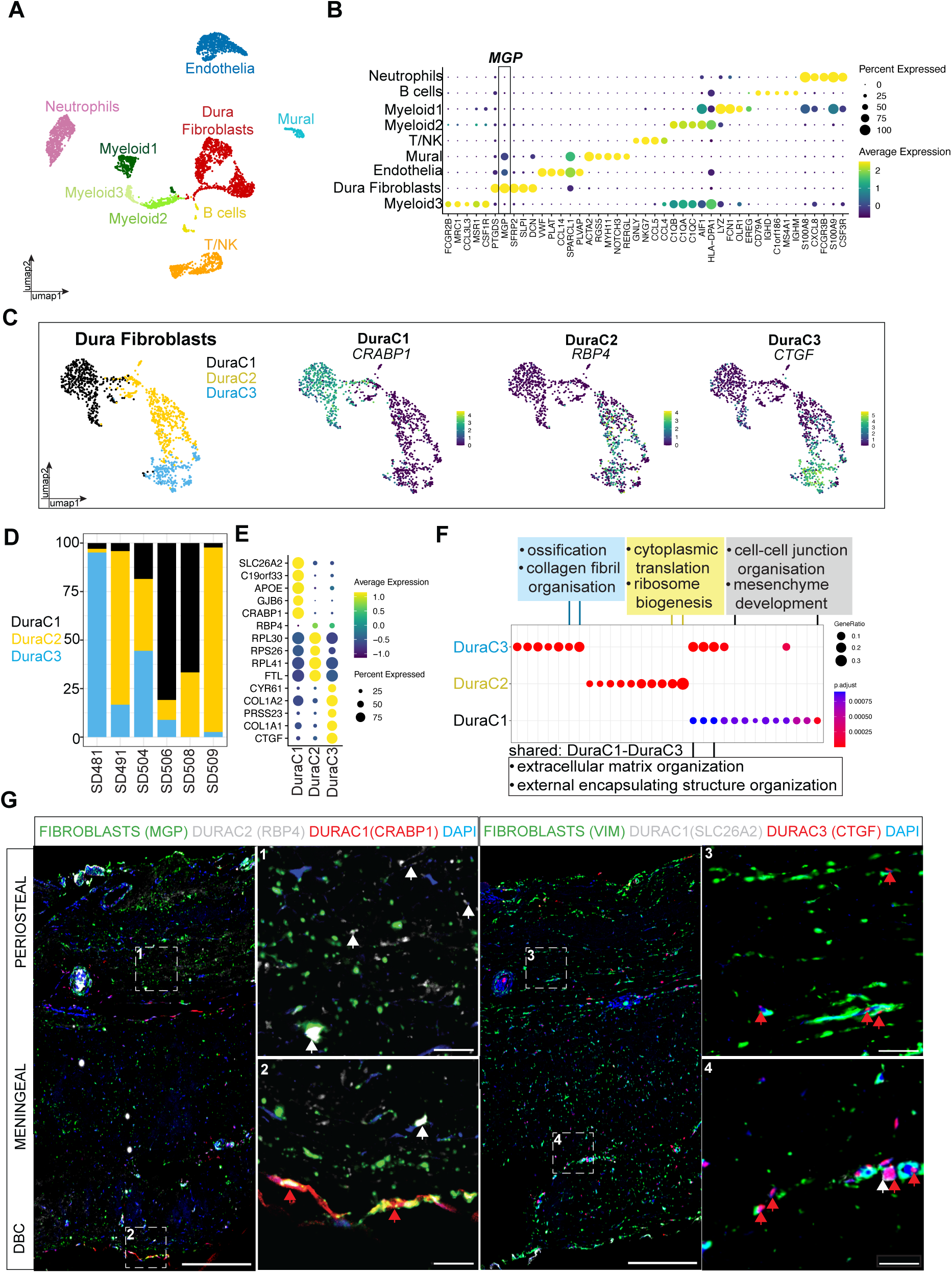
Characterisation and spatial localisation of dura fibroblast subpopulations. **(A)** UMAP of cell populations captured in the human dura: neutrophils; B cells; Myeloid1 cells; Myeloid2 cells; Myeloid3 cells; T/NK cells; mural cells; endothelia; and dura fibroblasts. **(B)** Dotplot of gene markers for each dura cell population. *MGP* is a key gene discriminating dura fibroblasts from other resident dura cells. Dot colour represents average expression level, and dot size represents the percentage of cells expressing the gene. **(C)** UMAP demonstrating the three primary fibroblast subpopulations in dura mater: DuraC1 (black), DuraC2 (yellow), and DuraC3 (blue). Adjacent feature plots show average expression of one key marker gene per dura fibroblast subpopulation. **(D)** Cell type proportions of each dura fibroblast subpopulation from 6 samples. **(E)** Dotplot of marker genes in each dura fibroblast subpopulation. Dot colour represents average expression level, and dot size represents the percentage of cells expressing the gene. **(F)** GO plot demonstrating representative biological pathways enriched in each dura fibroblast subpopulation: DuraC1 (cell-cell junction organization, mesenchyme development); DuraC2 (cytoplasmic translation, ribosome biogenesis); and DuraC3 (ossification, collagen fibril organisation). Select shared biological pathways are also shown: extracellular matrix organization and external encapsulating structure organization (DuraC1-DuraC3). Dot colour represents adjusted p-value, and dot size represents the gene ratio. **(G)** Protein immunolabelling showing spatial localisation of dura fibroblast subpopulations *in situ*: CRABP1 (red - DuraC1); RBP4 (white - DuraC2); and CTGF (white - DuraC3). Dashed boxes indicate the location of magnified regions. Scale bars = 100μm (low magnification images), 20μm (high magnification images).

Using differential gene expression analysis, we identified *MGP*, *SFRP2*, and *PTN* as markers of dura fibroblasts, compared to other dura microenvironment cells (**Figure 3B**). Endothelial cells showed high expression of *VWF* and *IFI27*, mural cells were marked by *ACTA2* and *NOTCH3* (see additional vascular cell type markers in **Supp. Figure 2**). We also identified several immune cell types in the dura mater which were distinguished by unique expression of the following genes: T/NK cells (*NKG7* and *CCL5*); B cells (*CD79A* and *IGHD*); neutrophils (*S100A8* and *S100A9*); and three myeloid cell subtypes (Myeloid cells1 [*LYZ* and *FCN1*]; myeloid cells2 [*C1QA* and *C1QB*] and myeloid cells3 [*MSR1* and *MRC1*]) (**Figure 3B**).

We then subset dura fibroblasts and identified three fibroblast subtypes (DuraC1, DuraC2, DuraC3) (**Figure 3C**). Five of the six samples contained three dura fibroblast populations; sample SD508 lacked DuraC3 cells (**Figure 3D**). DuraC1 was defined by high expression of *SLC26A2*, *GJB6*, and *CRABP1* (**Figure 3E**). Within the dura mater, the dura border cell (DBC) layer is a specialized substructure with barrier properties consisting of loosely arranged fibroblasts with gap junctions and/or desmosomes, solute carrier expression, and no collagen enrichment^19^. Accordingly, expression of *SLC26A2*, *GJB6*, and *CRABP1* (**Figure 3E**) led us to posit that DuraC1 fibroblasts comprise the DBC layer. Gene ontology analysis of overrepresented pathways in DuraC1 fibroblasts compared to all other dura fibroblast subtypes confirmed upregulation of biological pathways related to cell-cell junction organisation and mesenchyme development (**Figure 3F; Supp. Table 1**). Localisation of DuraC1 fibroblasts to the DBC layer was confirmed with protein labelling using MGP and CRABP1 (**Figure 3G**).

DuraC2 fibroblasts were differentiated from DuraC1 and DuraC3 fibroblasts by higher expression of *RPS26*, *RPL41*, *FTL*, and *RBP4* (**Figure 3E**). Gene ontology analysis of DuraC2 cells revealed upregulation of pathways related to cytoplasmic translation and ribosome biogenesis (**Figure 3F; Supp. Table 1**). Together, *RBP4* and *CRABP1* upregulation in DuraC2 and DuraC1 cells, respectively, suggests an overall upregulation of retinoic acid-associated pathways in human dura mater. Protein labelling showed fibroblasts expressing RBP4 are found throughout the meningeal and periosteal layers of the dura (**Figure 3G**).

DuraC3 cells were defined by expression of *CTGF*, *COL1A1*, and *BGN* (**Figure 3E**), genes associated with ossification, extracellular matrix, and collagen fibril organization; these pathways were strongly increased in DuraC3 compared to other dura fibroblast clusters (**Figure 3F; Supp. Table 1**). This suggests that DuraC3 fibroblasts reside in the periosteal and/or meningeal layers of the dura mater. Protein expression confirmed that DuraC3 collagen 1-expressing cells were strongly enriched in the periosteal layer of the dura, and to a lesser extent in the meningeal dura (**Figure 3G**).

In summary, DuraC1 fibroblasts match descriptions of DBCs, whereas DuraC2 and DuraC3 fibroblasts comprise the meningeal and periosteal dura layers, with DuraC3 fibroblasts being enriched in the periosteal layer.

### Characterisation and spatial localisation of arachnoid fibroblast subpopulations

Analysis of arachnoid mater from three samples revealed four primary cell populations: arachnoid fibroblasts; T/NK cells; neutrophils; and two myeloid cell populations (myeloid cells1, myeloid cells2) (**Figure 4A and B**). All cell types except neutrophils were observed in each sample. Neutrophils were not captured in only one sample (**Supp. Figure 3B**).

**Figure 4:**
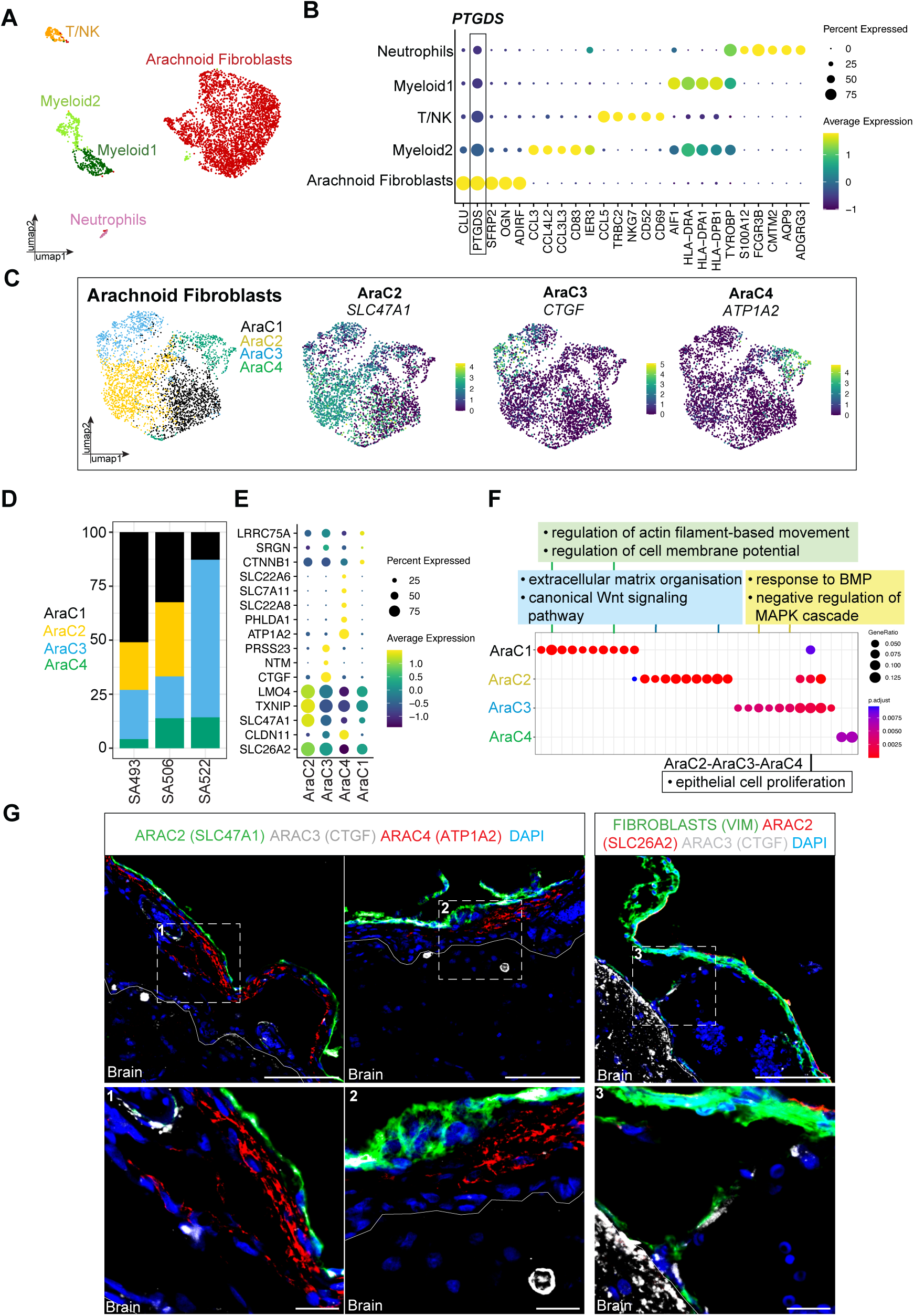
Characterisation and spatial localisation of arachnoid fibroblast subpopulations. **(A)** UMAP of cell populations captured in arachnoid: neutrophils; myeloid cells1; T/NK cells; myeloid cells2; and arachnoid cells. **(B)** Dotplot of gene markers for each arachnoid cell population. Dot colour represents average expression level, and dot size represents the percentage of cells expressing the gene. **(C)** UMAP demonstrating the four primary arachnoid fibroblast subpopulations: AraC1 (black); AraC2 (yellow); AraC3 (blue); and AraC4 (green). Adjacent feature plots showing expression of marker genes for each arachnoid fibroblast subpopulation. **(D)** Cell type proportions of each arachnoid fibroblast subpopulation from 3 samples. **(E)** Dotplot of marker genes of each arachnoid fibroblast subpopulation. Dot colour represents average expression level, and dot size represents the percentage of cells expressing the gene. **(F)** GO plot demonstrating representative biological pathways enriched in each arachnoid fibroblast subpopulation: AraC2 (response to BMP, negative regulation of MAPK cascade); AraC3 (extracellular matrix organization, canonical Wnt signaling); and AraC4 (regulation of actin filament-based movement, regulation of cell membrane potential). Select shared biological pathways are also shown: actin-mediated cell contraction (AraC3-AraC4); epithelial cell proliferation (AraC2-AraC3-AraC4); and ossification (AraC2-AraC3). Dot colour represents adjusted p-value, and dot size represents the gene ratio. **(G)** Protein immunolabelling showing spatial localisation of arachnoid fibroblast subpopulations *in situ*: SLC47A1 (green - AraC2); CTGF (white - AraC3); and ATP1A2 (red; AraC4). Dashed boxes indicate the location of magnified regions. Scale bars = 100μm (low magnification images), 20μm (high magnification images).

Arachnoid fibroblasts were distinguished from other arachnoid microenvironment cells by high expression of *PTGDS*, *CLU*, *SFRP2*, *OGN*, and *ADIRF* (**Figure 4B**). All arachnoid myeloid cells expressed *AIF1* and major histocompatibility complex II (MHC II) genes including *HLA-DRA* (**Figure 4B**). Myeloid cells2 uniquely expressed *CD83* (**Figure 4B**), an immunoglobulin expressed on antigen presenting cells, and *MS4A7* (**Figure 4B**), a marker for border-associated macrophages. Arachnoid-resident T cells and NK cells clustered together, expressing well-described markers including *CD3E* and *CD2*, and *CD52*, respectively (**Figure 4B**). Neutrophils showed unique expression of *S100A12* (**Figure 4B**).

To determine if arachnoid fibroblasts showed transcriptomic diversity, we subset them from the larger arachnoid dataset and identified four cellular subtypes (AraC1, AraC2, AraC3, AraC4) (**Figure 4C and D**). AraC2, AraC3, and AraC4 cells could be distinguished based on key differentially expressed genes. AraC2 fibroblasts upregulated solute carrier genes including *SLC26A7* and *SLC47A1* (**Figure 4E**). Enriched solute carrier genes suggests that AraC2 cells are important gatekeepers controlling molecule transport, including drugs and toxins, across the blood-CSF-barrier^20^. GO term analysis of AraC2 cells compared to all other arachnoid fibroblasts showed enrichment of pathways related to bone morphogenic protein and negative regulation of the MAPK cascade, which are important for cell growth and development (**Figure 4F; Supp. Table 1**). Cell localisation using protein labelling revealed that AraC2 cells were localised directly adjacent to the dura, suggesting that they comprise the outer layer of the arachnoid barrier cell layer (**Figure 4G**).

AraC3 fibroblasts showed highly unique expression of *COL1A1* and *CTGF* (**Figure 4E**) and over-representation of ECM production/organization pathways compared to other arachnoid fibroblast subtypes (**Figure 4F; Supp. Table 1**). Since arachnoid trabeculae cells are known to be uniquely bolstered by a network of ECM for stability/strength^21^, we suggest that AraC3 fibroblasts are arachnoid trabeculae cells. On occasion, these trabecular fibroblasts likely interact with pia fibroblasts, as we demonstrated with MERFISH (**Figure 2C, panel C2**). We confirmed that the CTGF protein marks arachnoid trabeculae cells (**Figure 4G**).

AraC4 fibroblasts were uniquely enriched with a variety of genes including *ATP1A2*, *NOV*, *CLDN11,* and *APOD* (**Figure 4E**). *CLDN11* is a tight junction molecule that has been described as an arachnoid barrier layer marker, suggesting that AraC4 cells may also form the arachnoid barrier layer along with AraC2 cells^22^. We also identified upregulation of biological pathways associated with cell membrane potential in AraC4 cells (**Figure 4F; Supp. Table 1**), confirming the utility of *ATP1A2* as a marker (**Figure 4E**). Protein labelling confirmed that ATP1A2-positive cells were widely expressed in the arachnoid barrier layer, mostly in fibroblasts in the central and inner rows facing the SAS (**Figure 4G**).

Overall, we identified markers for each arachnoid fibroblast subpopulation, with the exception of AraC1 cells. AraC2 cells localised to the outer arachnoid barrier layer, AraC4 cells to the inner arachnoid barrier layer, and AraC3 cells to the arachnoid trabeculae.

### Characterisation and spatial localisation of pia fibroblast subpopulations

Analysis of pia mater from three samples revealed five primary cell populations: pia fibroblasts; two endothelial cell populations (Endothelia1, Endothelia2); T/NK cells; myeloid cells; and mural cells (**Figure 5A and B**). Cell type proportions were estimated by subsampling one thousand cells per sample to avoid bias through sampling differences (**Supp. Figure 3C**).

**Figure 5:**
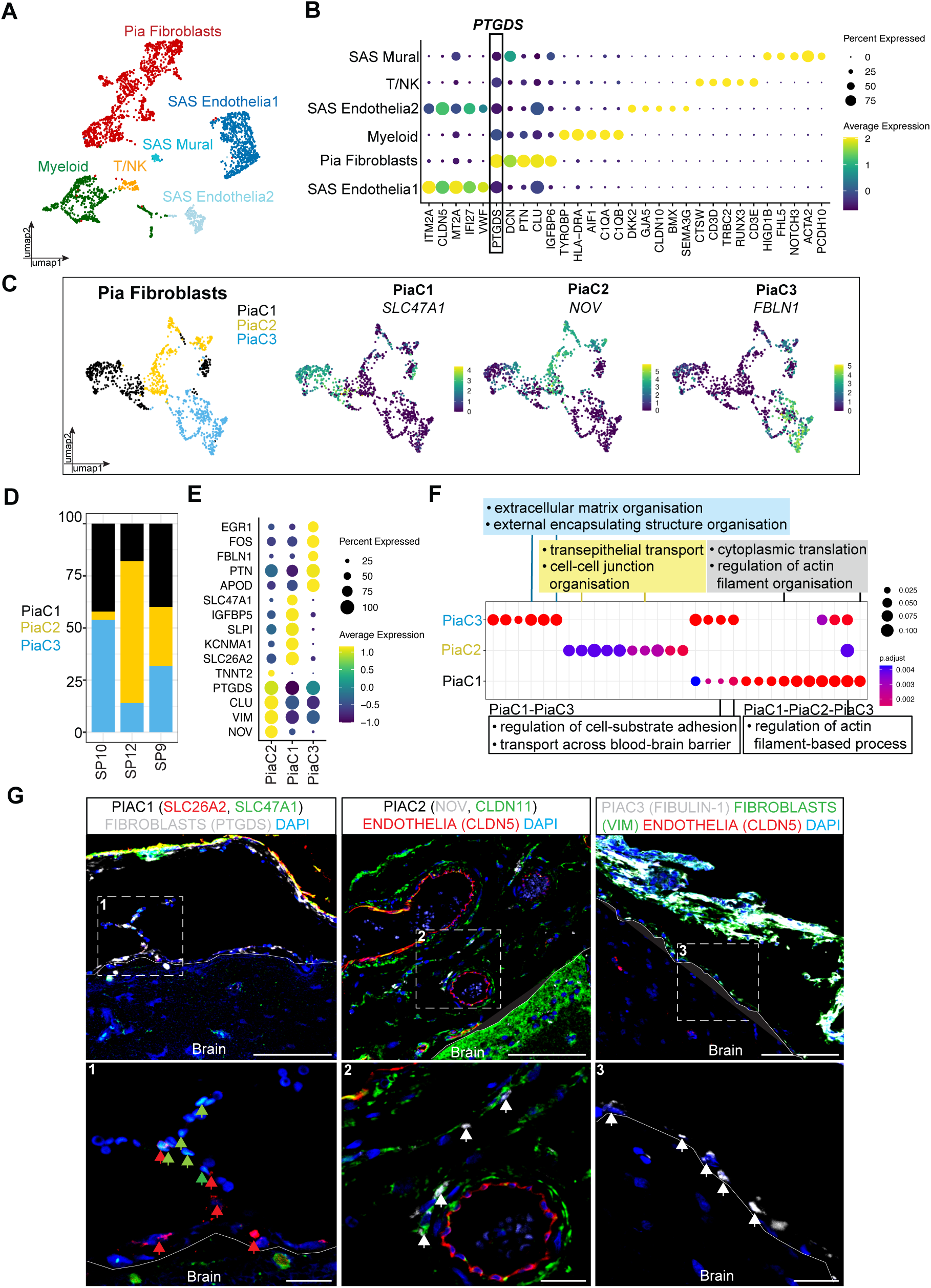
Characterisation and spatial localisation of pia fibroblast subpopulations. **(A)** UMAP of cell populations captured in the human pia: mural cells; T/NK cells; endothelia2, myeloid cells; pia cells; and endothelia1. **(B)** Dotplot of marker genes for each pia cell population. Dot colour represents average expression level, and dot size represents the percentage of cells expressing the gene. **(C)** UMAP demonstrating the three primary pia fibroblast subpopulations: PiaC1 (black); PiaC2 (yellow); and PiaC3 (blue). Adjacent feature plots showing expression of marker genes for each pia fibroblast subpopulation. **(D)** Cell type proportions of each pia fibroblast subpopulation from 3 samples. **(E)** Dotplot of marker genes of each pia fibroblast subpopulation. Dot colour represents average expression level, and dot size represents the percentage of cells expressing the gene. **(F)** GO plot demonstrating representative biological pathways enriched in each pia fibroblast subpopulation: PiaC1 (cytoplasmic translation, regulation of actin filament organization); PiaC2 (transepithelial transport, cell-cell junction organization); and PiaC3 (extracellular matrix organization, external encapsulating structure organization). Select shared biological pathways are also shown: regulation of cell-substrate adhesion and transport across blood-brain barrier (PiaC1-PiaC3); and regulation of actin filament-based process (PiaC1-PiaC2-PiaC3). Dot colour represents adjusted p-value, and dot size represents the gene ratio. **(G)** Protein immunolabelling showing spatial localisation of pia fibroblast subpopulations in *in situ*: SLC47A1 (green - PiaC1); SLC26A2 (red - PiaC1); PTGDS (white - PiaC2); Fibulin-1 (green - PiaC3); and APOD (red; PiaC3). Dashed white boxes indicate the location of magnified regions. Scale bars = 100μm (low magnification images), 20μm (high magnification images).

Pia fibroblasts were distinguished from other microenvironment cells by upregulation of the leptomeningeal markers *PTGDS*^23^, *CLU*, *DCN*, and *OGN* (**Figure 5B**). Both endothelial cell clusters expressed *CLDN5*. Endothelia1 cells were also enriched in *ITM2A*, *MT2A,* and *VWF* (**Figure 5B**), whereas Endothelia2 cells were enriched for *SEMA3G* (**Figure 5B**). The primary immune cell populations were T/NK cells (*CD3D*, *CD2*) and myeloid cells (*AIF1*). The myeloid cell cluster also showed high expression of antigen presenting genes (*HLA-DRA*) and genes associated with inflammation initiation (*C1QB*, *C1QA*). Lastly, we identified a *NOTCH3*-positive mural cell cluster (**Figure 5B**).

We next subset *PTGDS*-positive pia fibroblasts for further analysis. In doing so we identified three pia fibroblast subtypes (PiaC1, PiaC2, PiaC3) (**Figure 5C and D**). PiaC1 cells were specifically enriched in the solute carriers *SLC26A2* and *SLC47A1* compared to other pia fibroblasts (**Figure 5E**). The primary biological pathways enriched in PiaC1 cells were cytoplasmic translation and regulation of actin filament organisation (**Figure 5F; Supp. Table 1**). Regulation of cell-substrate adhesion and transport across the blood-brain barrier were shared functions among PiaC1 and PiaC3 cells (**Figure 5F; Supp. Table 1**). Protein labelling revealed that most PiaC1 cells are situated close to the brain surface (**Figure 5G**).

PiaC2 fibroblasts were enriched in the expression of *NOV*, a gene associated with the ECM and basement membrane formation^2,24,25^, and *CLDN11* (**Figure 5E**). Although well-described in arachnoid barrier cells^22,26^, *CLDN11* has not been previously linked to pia fibroblasts. Pathways predicted to be upregulated in PiaC2 cells included transepithelial transport and cell-cell junction organization (**Figure 5F; Supp. Table 1**). Using protein labelling, we observed NOV expression in a subset of pia fibroblasts wrapped around large blood vessels in the SAS (**Figure 5G**), suggesting that PiaC2 fibroblasts represent a subpopulation involved in regulating ion transport between the CSF and SAS blood vessels. We also suggest that PiaC2 cells, based on their unique SAS localisation, are the most likely pia fibroblast subtype to interact with arachnoid trabeculae AraC3 cells (**Figure 2C, panel C2**).

PiaC3 fibroblasts highly expressed *FBLN1* and *COL1A1* (**Figure 5E**), both of which are associated with ECM production and maintenance. Collagen fibres have been observed to be enriched in the brain-covering pia fibroblasts, some of which are closely connected to arachnoid trabeculae^27^. Fibulin-1 protein labelling revealed that PiaC3 fibroblasts, like PiaC1 fibroblasts, reside primarily at the base of the pia membrane directly adjacent to the brain; however, some were also detected around blood vessels in the SAS (**Figure 5G**).

In summary, we demonstrate that PiaC1 and PiaC3 fibroblasts cells are likely brain-covering pia cells, whereas PiaC2 fibroblast cells are SAS blood vessel-covering cells.

### Cell-cell interaction analysis uncovers VEGF- and IGF-mediated crosstalk between fibroblasts and endothelial cells of the dura and subarachnoid space (SAS)

Having created a transcriptomic, functional, and spatially-resolved cellular map of dura, arachnoid, and pia fibroblast subtypes within each layer of the human meninges, we used it to determine crosstalk between meningeal fibroblasts and other cells in their microenvironment. We focused on fibroblast interactions with endothelial cells in the dura and the SAS because the relationship between vasculature and their surrounding meningeal fibroblasts in these two unique meningeal microenvironments has not been well characterised^28^.

We first completed an unbiased prediction of cell-cell interactions between all cells comprising the dura mater (**Figure 6A**; **Supp. Figure 8A**). This analysis showed that endothelial cells have the highest number of predicted ligand-receptor (LR) pairs with other cell types (nine hundred and ninety-four LR pairs; p<0.05) (**Supp. Figure 8A**). DuraC1 fibroblasts (nine hundred and ninety-four LR pairs; p<0.05) and Myeloid cells1 (nine-hundred and sixty-five LR pairs; p<0.05) also showed high numbers of predicted interaction pairs (**Supp. Figure 8A**). Nearly all dura cell types expressed vascular endothelial growth factor (VEGF) ligands that bind to their respective receptors on dura endothelial cells (**Figure 6A**). In particular, mast cells and myeloid cells1 possessed the highest interaction scores for *VEGFA*-*FLT1* interactions, demonstrating a role for myeloid cell-endothelial cell interactions in the maintenance of dura vasculature.

**Figure 6:**
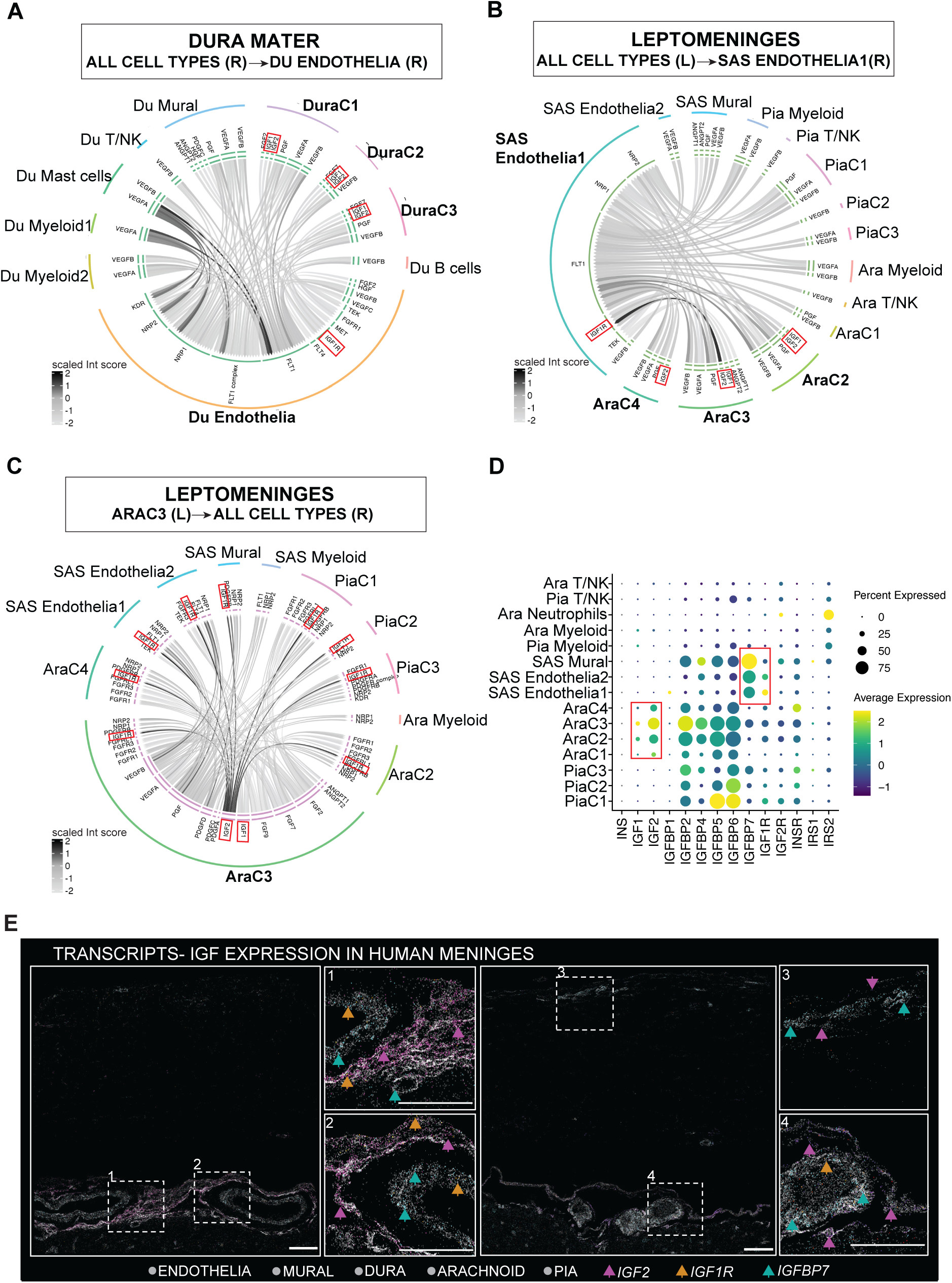
Cell-cell interaction analysis predicts VEGF and IGF-mediated crosstalk between fibroblasts and endothelial cells of the dura and subarachnoid space. **(A)** Circle plot demonstrating the top significantly enriched ligand-receptor interactions pairs between all cell types in the dura and dura endothelial cells. Line colour indicates scaled interaction score. **(B)** Circle plot demonstrating the top significantly enriched ligand-receptor interactions pairs between all cell types in the leptomeninges and SAS Endothelia1. Line colour indicates scaled interaction score. **(C)** Circle plot demonstrating the top significantly enriched ligand-receptor interactions pairs between AraC3 fibroblasts and all other cell types in the leptomeninges. Line colour indicates scaled interaction score. **(D)** Dot plot of canonical insulin growth factor signaling gene expression in all leptomeningeal cell types. Dot colour indicates average expression and dot size indicates percentage of cells expressing the gene. **(E)** Two representative MERFISH panels from 2 samples demonstrating expression of *IGF2* (purple), *IGF1R* (orange), and *IGFBP7* (blue) transcripts in the adult human meninges. Dotted white boxes (E1-4) indicate magnified regions. Scale bars = 100μm.

**Figure 7:**
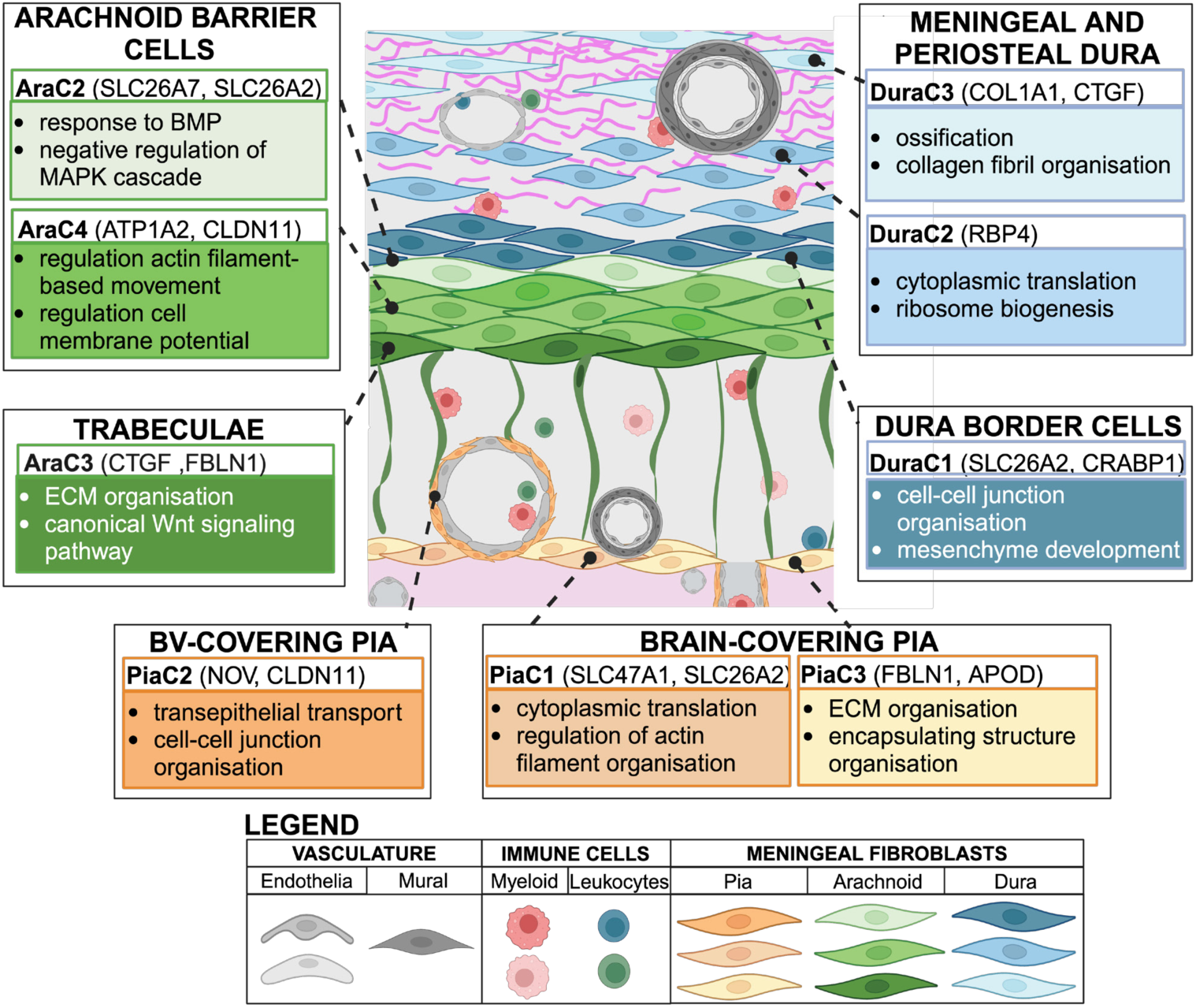
Spatial arrangement of layer-specific fibroblast subpopulations in the human meninges. This schematic provides a proposed neuroanatomical arrangement of the fibroblast subtypes identified in each layer of the human meninges by scRNAseq and confirmatory protein staining (Figures 3-5). DuraC1 (*SLC26A2*, *CRABP1*) fibroblasts likely represent dura border cells, whereas DuraC2 (*RBP4*) fibroblasts appear to be scattered throughout the meningeal and periosteal dura. DuraC3 (*COL1A1*, *CTGF*) fibroblasts may be enriched in the periosteal dura, especially given unique predicted interactions with dura endothelial cells that reside specifically in the periosteal layer (Figure 6). AraC1 cells were not transcriptomically distinct and were therefore not included in our meningeal fibroblast schematic. AraC2 (*SLC26A7*, *SLC26A2*) and AraC4 (*ATP1A2*, *CLDN11*) fibroblasts appear to represent outer and inner layers of arachnoid barrier cells, respectively, whereas we suggest AraC3 (*CTGF*, *FBLN1*) fibroblasts correspond to arachnoid trabeculae. Although we had limited instances in which to confirm localisation of AraC3 cells due to few intact trabeculae in our tissue microarray, this hypothesis is further supported by unique predicted interactions between AraC3 cells and SAS Endothelia1 (Figure 6). Finally, both PiaC1 (*SLC47A1*, *SLC26A2*) and PiaC3 (*FBLN1*, *APOD*) fibroblasts were typically localised directly adjacent to the brain, whereas PiaC2 (*NOV*, *CLDN11*) fibroblasts more often enshrouded vessels in the subarachnoid space. Together, these data provide the first complete layer-specific reconstruction of cellular heterogeneity in the human meninges.

Dura cells were also predicted to release insulin growth factor (IGF) ligands, including *IGF1* and *IGF2*, that bind to *IGF1* receptors on endothelial cells (**Figure 6A**). We further investigated the expression levels of molecules involved in the IGF1 signaling cascade and observed unique expression of IGF ligands (primarily *IGF2*) by dura fibroblasts, especially DuraC3 cells (**Figure 6B; Supp. Figure 8B**). We showed earlier that DuraC3 cells are located primarily in the meningeal and periosteal dura (**Figure 3G**). Given dura blood vessels are located primarily in the periosteal dura, or between the meningeal and periosteal layers, this predicted interaction between DuraC3 cells and endothelial cells further supports the finding that DuraC3 fibroblasts are periosteal layer resident cells. Genes coding for other key proteins involved in IGF signaling, such as *IGFBP2/4/5/6/7*, were also expressed primarily by dura fibroblasts, but also endothelial cells and mural cells (**Figure 6A; Supp. Figure 8B**).

Having predicted *VEGF* and *IGF* signaling between dura fibroblasts and endothelial cells, we assessed if similar pathways were regulated by these two cell types in the leptomeninges. To do this, we merged the pia and arachnoid datasets to assess the SAS vasculature and its surrounding fibroblasts and immune cells (**Supp. Figure 8C**). Like in the dura, we completed an unbiased prediction of cell-cell interactions between all cells comprising the leptomeninges (**Figure 6B**). Of all the cell types profiled in the integrated leptomeningeal object, AraC3 was predicted to have the highest number of interactions with other cell types (one thousand and fifty-eight L-R pairs; p<0.05) (**Supp. Figure 8D**). We previously showed that AraC3 fibroblasts represent a unique subtype confined to arachnoid trabeculae (**Figure 4G**); these interaction analysis data further support this finding given arachnoid trabeculae traverse the entire SAS interacting with other fibroblasts, vascular cells, and immune cells^29^. All leptomeningeal fibroblasts expressed VEGF ligands that bind to their respective receptors on SAS endothelial cells1 and 2 (**Figure 6B; Supp. Figure 8E**). In particular, AraC2 and AraC3 fibroblasts possessed the highest interaction scores for *VEGFB*-*FLT1* interactions.

In addition to *VEGF*s, AraC3 fibroblasts also expressed angiopoietins (*ANGPT1/2*), which were predicted to bind to TEK tyrosine kinase (*TIE2*) on SAS endothelia cells1 (**Figure 6B**). *IGF1* and *IGF2* were significantly expressed by all three arachnoid fibroblast subtypes (AraC2, AraC3, AraC4), but not by pia fibroblasts or immune cells in the SAS (**Figure 6B**). Interestingly, by visualizing all significantly enriched ligands expressed by AraC3 (**Figure 6C**), we uncovered a strong interaction between AraC3- produced *IGF2* and *IGF1R* on all leptomeningeal fibroblast subpopulations and vascular cells (**Figure 6C and D; Supp. Figure 8E**). Finally, IGFBPs were also expressed by a variety of leptomeningeal cell types in the SAS, including pia and arachnoid fibroblasts and vascular cells (**Figure 6D**). In particular, *IGFBP7*^30^ was highly expressed in vascular cells of the SAS (**Figure 6D**). MERFISH confirmed expression of *IGF2* in meningeal cells, whereas *IGF1R* and *IGFBP7* were primarily expressed in vascular cells of all layers of the meninges as predicted by the interaction analysis (**Figure 6E**).

Together, these representative interaction analyses demonstrate key pathways relevant to the communication of layer-specific human meningeal fibroblasts and vascular cells.

## Discussion

These data establish the first layer-resolved atlas of the adult human meninges. Using scRNAseq, MERFISH, and protein labelling, we confirm the spatial localisation of all meningeal cell types, decode ten distinct fibroblast cell populations, and pinpoint their layer-specific locations within distinct neuroanatomical niches. We then predict how meningeal fibroblasts interact with vascular cells in their immediate environment and unveil the involvement of IGF2-IGF1R signaling in this relationship. Together, this atlas provides a comprehensive resource to support future investigations of human meningeal architecture, cell-cell interactions underpinning homeostatic meningeal function, and the normotypic context to understand alterations to the meningeal microenvironment that occur in CNS diseases.

The dura mater is known to be strongly enriched in extracellular matrix components and is subdivided into a DBC layer, a meningeal dura layer, and a periosteal dura layer^31^. DuraC1 fibroblasts localised to the innermost layer of the dura and highly expressed genes associated with junctional protein organization pathways (i.e., *GJB6*) and solute carriers (i.e., *SLC26A2*). In adult mice, solute carrier genes are known to be enriched in the DBC layer^13^. In humans, the DBC is known to consist of flattened fibroblasts with prominent intercellular junctions and minimal collagen in the surrounding extracellular space^31^. These data highly suggest that DuraC1 fibroblasts are DBC layer fibroblasts.

DuraC3 fibroblasts localised to the periosteal and meningeal layers of the dura and showed enrichment of genes associated with ossification and collagen fibril organization. Cell-cell interaction analysis predicted that DuraC3 cells interact with dural vasculature through molecules such as IGFs, further strengthening the finding that they reside in the periosteal and/or meningeal layers. Finally, localisation of DuraC2 fibroblasts also demonstrated that these cells were located in the periosteal and/or meningeal layers. Previous studies have suggested that retinoic acid pathway regulation is relevant for meninges-instructed brain development^32^, but how different dura fibroblast subpopulations might play a role is unknown.

The arachnoid mater is subdivided into arachnoid trabeculae, which span the entire SAS anchoring the pia and SAS vasculature to the arachnoid, and the arachnoid barrier cell (ABC) layer, which abuts the DBC layer. AraC3 fibroblasts localised to arachnoid trabeculae and were enriched in collagens and other ECM molecules (i.e., *FBLN1*) compared to other arachnoid fibroblast subpopulations. Trabecular fibroblasts are known to be supported by extracellular collagen to help withstand the pressure exerted by CSF circulating in the SAS^33^. Cell-cell interaction analysis also predicted IGF2-mediated interactions of AraC3 cells with both pia fibroblasts and SAS vascular cells. Given these data, it is highly likely that AraC3 fibroblasts are arachnoid trabeculae fibroblasts.

AraC4 fibroblasts localised to the ABC layer. In mice, the ABC layer consists of several densely packed layers of CD166-positive fibroblasts^26^. ABC fibroblasts are connected by tight junctions such as CLDN11 as previously demonstrated in mice and human fetuses^22,26,34^. We also identity another population of fibroblasts, AraC2 fibroblasts, that localise to the ABC, between AraC4 fibroblasts and the DBC layer. This finding suggests that there are two distinct human ABC fibroblast subpopulations.

The pia mater is formed by two distinct layers^13,29^. Epi-pia fibroblasts, which wrap around blood vessels in the SAS^35^, and brain-covering pia fibroblasts, which secrete a thin basal lamina composed of densely arranged ECM fibers^24^. PiaC2 fibroblasts localised to vessels in the SAS and upregulated genes associated with cell-cell junction regulation, including *CLDN11*ma tight junction protein commonly described in the ABC^26^. Cell-cell interaction analysis also predicted IGF- and VEGF-mediated crosstalk between PiaC2 fibroblasts, vascular cells of the SAS, and AraC3 fibroblasts, which we localised to arachnoid trabeculae. These data highly suggest that PiaC2 fibroblasts are blood vessel-covering pia fibroblasts. PiaC3 fibroblasts localised to the cortical surface and were enriched in genes associated with ECM organization compared to other pia fibroblasts subpopulations. This confirmed that PiaC3 fibroblasts are brain-covering pia fibroblasts.

Interpretation of atlas data should be made with knowledge of important experimental limitations. Due to surgical constraints, meninges samples were sourced from skull convexity regions only. As a result, there is limited representation of all potential cell states from other regions of the brain, including a lack of representation of arachnoid granulation-associated cells. This limitation will likely be most relevant when examining region-specific pathologies such as meningiomas, which derive from the meninges covering different brain regions. Lastly, we did not capture any lymphatic endothelial cells in our samples. This could be due to our chosen scRNAseq method, or because we did not sequence multiple meningeal regions in each sample, which would increase the chance of capturing a significant number of lymphatic vessels.

In summary, we developed the first complete layer-resolved atlas of the adult human meninges by integrating scRNAseq, MERFISH, and immunolabelling. Beyond localising all primary meningeal cell types within each layer, we critically dissect meningeal fibroblasts and discover ten unique subpopulations. Future studies could validate the predicted functions of layer-specific fibroblast subpopulations, including DBC fibroblasts, ABC fibroblasts, and both blood vessel- and brain-covering pia fibroblasts. These data can be further analyzed using an [open science platform](https://singlocell.openscience.mcgill.ca/).

## Method

### Human tissue samples

Surgeries were performed and meninges samples were harvested under a protocol approved by the Montreal Neurological Institute-Hospital research ethics board. Consent was given by all patients. Pre- operative magnetic resonance imaging was performed for surgical planning. Six dura mater samples, three arachnoid mater samples, and three pia mater samples were acquired from regions adjacent to, but still far from, the tumor site (Table 1).

### Tissue processing for single cell RNA sequencing

All meningeal samples were placed immediately on ice following surgical extraction and were processed within 30 minutes (min). Dura and arachnoid mater samples were washed three times using Wash Buffer (sterile phosphate buffered saline (PBS) with 1 % penicillin and streptomycin). Pia mater samples were washed using the same Washing Buffer with an additional 0.04 % bovine serum albumin (BSA). All specimens were minced into fragments of less than 1 mm in size. Dura and arachnoid samples were digested using collagenase, and pia samples in a 0.25 % trypsin solution. Both solutions contained DNAse (50 U/ ml) and MgCl_2_ and were digested for 1-2 hours at 37 ^◦^C. Digested specimens were washed three times with Wash Buffer, and large debris were removed with a 70 μm strainer. Red blood cells were removed using a density gradient in a 1:1 volume ratio with the sample (Lymphoprep, Axis-Shield). Cell pellets were resuspended in PBS with 0.04 % BSA to a final concentration of 1000 cells/µl.

### Single cell RNA sequencing – Drop-seq protocol

The Drop-Seq protocol was performed by 10X using the cell ranger pipeline. The sequencing protocol was adapted from Couturier et al., 2020^36^. In summary, layer-specific meningeal cell suspensions were generated from surgically extracted specimens and immediately sequenced. V2 or v3 Single Cell 3’ Reagent Kits (CG0052 10x Genomics) were used. RNA libraries were generated using the GemCode Single-Cell Instrument (10x Genomics, Pleasanton, CA, USA). For library generation, single cells, barcoded beads and reagents for sample preparation were sorted into aqueous droplets. The barcoded beads contained a 30 bp oligo(dT) bound to the mRNA, 8bp molecular index that was unique for each mRNA strand, 12bp barcode unique to the cell, and a barcode universal for all beads (Single Cell 3’ Library, Gel Bead Kit v2/v3, Chip Kit (P/N 120236 P/N 120237 10x Genomics). Single cells were lysed within the droplet and captured on the barcoded bead. mRNA strands were reverse transcribed into cDNA and amplified. Sequencing ready libraries were purified with SPRIselect beads, controlled for yield and sized distribution (LabChip GX Perkin-Elmer), and quantified using qPCR (KAPA Biosystems Library Quantification Kit for Illumina platforms P/N KK4824). Demultiplexing of cell barcodes and unique molecular identifiers was followed by alignment of single end reads to a reference genome (GRCh38, Cell Ranger Pipeline).

### Single cell RNA sequencing data analysis

Single cell RNA sequencing data analysis steps were primarily performed using Seurat V4^37^. Quality control steps differentiated technical noise from real biological variation after sequencing. Cells with high mitochondrial gene content (> 5 %) and cells with less than 200 or more than 2500 features were removed. Count data were scaled by linear transformation. Linear dimensionality reduction of the 2000 selected most variable features per dataset was applied. Principal component analysis (PCA) was performed on scaled data with variable features. Elbow plot has been used to determine the number of most representative components. K-nearest neighbor (KNN) was applied based on euclidean distance in the principal component space. Louvain algorithm was used to cluster cells based on transcriptomic similarity. Resolution parameters of clustering were tested between values of 0.1-1 to identify the most ideal clustering to represent cell types present. non-linear reduction technique Uniform Manifold Approximation and Projection (UMAP) was used to visualize clusters in 2D for data exploration. Differential gene expression was performed to analyze the expression of markers that are most differential for cells of one cluster compared to cells in other clusters. Cell types were annotated by profiling expression of known canonical marker genes from literature.

Downstream gene ontology (GoTerm) analysis was predicted through Clusterprofiler (v4.12.6)^38^. Top 500 significantly differentially expressed genes (p < 0.05) of each clusters will be associated with functional terms described on existing databases including ReactomePA, Go.db, enrichplot and DOSE and visualised^38^. Cell-cell interaction analysis was performed using CellPhoneDB (v2.1.7)^39^. CellPhoneDB is a repository of cell interactions based on already published datasets. It predicts receptors and ligands of cell clusters that may interact with one another. Counts and metadata of the RNA assay were extracted from the Seurat object and significant interaction scores (p < 0.05) and likelihood of interaction among annotated cell clusters determined after pipeline developed by Efremova et al, 2020. Predicted interactions were visualised using the interactive InterCellar webpage^40^.

### Copy number variation analysis

To identify potential large-scale chromosomal alterations that were affecting large chromosomal areas, we were inferring copy number variations (CNVs) from single cell RNA sequencing data (inferCNV i6, Trinity CTAT Project, Broad Institute). Randomly sampled immune cells were used as reference cells to define the baseline of gene expression. Gene expression levels of adjacent genomic regions were averaged to reduce noise and variation to determine gene expression levels. Due to sensitivity limits, just large genomic areas with many CNV events compared to low CNV numbers of adjacent genome areas were detectable. The cutoff value is set to 0.1 which determines which genes were used for the analysis. CNV calling threshold was set to > 40 % of each chromosome with Wilcoxon test p-value < 0.001. Red is visualizing overexpression or amplification, and blue, chromosome loss or reduced gene expression of large chromosomal regions. The rows of the heatmap correspond to cells ordered by hierarchical clustering by sample.

The columns display genes ordered by chromosome position (inferCNV i6 of the Trinity CTAT Project, Broad Institute; 10X Genomics)^36,41,42^. Downregulation of chromosome 6 in the fibroblast cells of pia, arachnoid, and dura samples is to be expected since referenced immune cells show a naturally strong upregulation of MHC II related genes located on chromosome 6.

### Tissue processing and microarray preparation for immunohistochemistry

Immediately after surgical extraction, leptomeningeal and/or dura tissue samples were rinsed with RBC lysis buffer before drop-fixation in 10 % formalin for 24 h followed by several rinses with PBS. Dura had to be extracted separately from the brain and overlying leptomeninges, which led to partial disruption of the adjoining dura border cells and arachnoid barrier cells (Figure 1B; Supp. Figure 1A). Fixed tissues were stored short term in 70 % ethanol overnight before tissue processing. For processing, samples were immersed in ethanol baths of increasing concentration (50 % −100 %) followed by xylene and xylene-paraffin baths to adequately dehydrate the tissue. Samples were then manually embedded in paraffin and sectioned (5µM) using a standard microtome (SLEE, CUT 6062) and mounted (Fisher, Superfrost Plus) using a water bath set at 60 degrees. After embedding and sectioning, the remainder of the blocks were used for creation of a tissue microarray for high-throughput H&E staining and protein labelling using a Tyramide protocol. In brief, the microarray was generated by taking two, 1 mm x 1 mm tissue cores (when possible) from each tissue block listed in Table 2 (dura and leptomeninges). Dura and leptomeninges cores were all embedded in a single paraffin block to create the microarray. Two cores of human tonsil tissue were added to the microarray as a positive control. 5 µm thick paraffin sections of the microarray were cut using a microtome (SLEE, CUT 6062) and mounted on glass slides (Fisher, Superfrost Plus) as described previously.

**Table 2.** Fresh human brain and meninges samples used for the tissue microarray.

| Sample | Tissue | Age | Sex | Pathology |
| --- | --- | --- | --- | --- |
| S536 | Brain, Leptomeninges, Dura | 69 | m | Metastatic Adenocarcinoma, Lung |
| S539 | Brain, Leptomeninges, Dura | 67 | m | Glioblastoma |
| S546 | Brain, Leptomeninges, Dura | 65 | f | Metastatic Non-Small Cell Carcinoma |
| S548 | Brain, Leptomeninges, Dura | 69 | m | Metastatic Adenocarcinoma |
| S551 | Brain, Leptomeninges, Dura | 79 | f | Metastatic Carcinoma |
| S535 | Dura | 76 | m | Glioblastoma |
| S540 | Dura | 73 | f | Metastatic Adenocarcinoma, Lung |
| S541 | Dura | 51 | f | Metastatic Carcinoma, Breast |
| S542 | Dura | 51 | m | Meningioma, Gr I |
| S552 | Dura | 67 | m | Glioblastoma |
| S606 | Brain, Leptomeninges, Dura | 72 | m | Glioblastoma |

### Hematoxylin & Eosin (H&E) staining

FFPE tissue slides were dewaxed with xylene (5 min; 2x 2 min), rehydrated in a series of ethanol solutions (2x 2 min at 100 %, 2x 2 min at 95 %, 1x 2 min at 70 %), and rinsed once for 5 min in ddH_2_O. Tissue sections were submerged in Hematoxylin 560 (Leica) for 3 min. Sections were washed using running water until clear (3-5 min) before being submerged in Define Buffer (Leica) for 45 s followed by a rinse with ddH_2_0 for 1 min. Slides were dipped into Bluing Buffer (Leica) for 30 s. Two quick washes with ddH_2_O for 15 s followed. Sections were dipped in 100 % ethanol for 10 seconds and incubated in Eosin Y Solution for 4 min. Lastly, tissue sections were rinsed using 100 % ethanol for 10 s and mounted using Gold Antifade mounting medium.

### Immunohistochemistry (Tyramide amplification)

Following deparaffinization as described above, the following steps were repeated for every antibody used: antigen retrieval with 10 % Citrate buffer (Sigma, C9999) in ddH2O in an antigen retrieval chamber (100 °C, 10 min), followed by cooling and washing 3x in PBS. Blocking was performed for 60 min at room temperature (RT) with a buffer containing 10 % Goat Serum (Invitrogen, B40923). Primary antibodies (Table 3) were diluted in 2 % BSA in PBS and incubated overnight at 4 °C. Subsequent steps were completed with Alexa Tyramide Super Boost Kits: B40913, B40923, B40922, B40912 (Thermo Fisher). After washing 3x for 10 min in PBS, slides were incubated with poly-HRP conjugated secondary antibodies for 60 min at RT and washed 3x with PBS. 100 µl of tyramide working solution was then added for 8 min at RT, followed by addition of 100 µl of stop reagent for 5 min. Finally, DAPI solution (1:1000 in PBS) was applied for 5 min followed by 3x 10 min washes in PBS. Gold Antifade Mounting Medium was used to mount all slides.

**Table 3.** Primary antibodies used for tissue microarray immunohistochemistry.

| Antibody | Species | Supplier | Product Number |
| --- | --- | --- | --- |
| APOD | rb | Thermo Fisher | PA5-27386 |
| AQP1 | rb | Thermo Fisher | MA532593 |
| AQP4 | ms | abcam | ab9512 |
| aSMA | rb | Thermo Fisher | 17H19L35 |
| ATP1A2 | rb | Thermo Fisher | 16836-1-AP |
| CD166 | rb | Thermo Fisher | PA5-12513 |
| CD31 | ms | abcam | ab9498 |
| CLDN5 | rb | abcam | ab131259 |
| CLDN11 | rb | Thermo Fisher | 36-4500 |
| CRABP1 | ms | Thermo Fisher | MA3-813 |
| CTGF | rb | abcam | ab6992 |
| Fibulin | ms | Thermo Fisher | MA5-24598 |
| IBA1 | rb | Thermo Fisher | PA5-27436 |
| MGP | rb | Thermo Fisher | 10734-1-AP |
| NOV | rb | Thermo Fisher | PA5-54055 |
| PTGDS | rb | Thermo Fisher | 10754-2-AP |
| RBP4 | rb | Thermo Fisher | 11774-1-AP |
| SLC26A2 | rb | Thermo Fisher | PA5-63524 |
| SLC47A1 | rb | Thermo Fisher | PA5-54541 |
| VIM | ms | Millipore | CBL202 |

### Confocal microscopy and image quantification

Fluorescent images were acquired using a Confocal Microscope (Zeiss LSM 700) with the Zen 2010 software at 20x. Images were converted to RGB images using ImageJ2 (version 2.14.0).

### Multiplexed error-robust fluorescence in situ hybridization (MERFISH) gene panel design

MERFISH is a spatial transcriptomic technique that allows for the detection of cell types *in situ* through simultaneous detection of hundreds of RNA molecules. We created a three hundred gene panel using the single cell RNA sequencing data described previously. We determined differentially expressed genes for cell types and pathways of interest with highest sensitivity (level of expression) and specificity (uniquely expressed in the population relative to all others). Ribosomal and mitochondrial genes were excluded from the list of genes if detected. To determine the number of fragments per kilobase of transcripts per million mapped reads (FPKM values) for our gene list, we performed bulk RNA sequencing on eight meningioma samples we previously performed single cell RNA sequencing on (see Table 1). Using the Vizgen Gene Panel Design Platform, genes with insufficient target regions were removed. Genes with high FPKM values were added to a subsequent panel, or removed, as leaving them in could lead to optical crowding. For the same reason, the total FPKM value of all genes in the panel was kept below nine thousand.

### MERFISH sample preparation and run

Samples were prepared according to the FFPE Sample Preparation User Guide provided by Vizgen. In short, the paraffin-embedded tissue microarray was sectioned, and the first ten sections were removed to avoid dry tissue with low RNA quality. Subsequent 5 µm sections of the tissue microarray were placed on MERSCOPE-specific slides and stored at –20 °C. Using the RNA Verification Kit, control probes targeting EEF2 were applied to examine RNA quality. Quality parameters including probe colocalisation, signal intensity, and frequency around DAPI+ nuclei were determined. If sufficient RNA quality was established, we applied the three hundred gene panel to the tissue microarray using the FFPE Sample Preparation Kit. The tissue sample underwent deparaffinization, RNA de-crosslinking and anchoring, gel embedding, tissue clearing, autofluorescent quenching, and gene panel hybridization. Lastly, DAPI and PolyT staining was applied before imaging on the MERSCOPE Instrument as according to the FFPE Sample Preparation Kit from Vizgen.

### MERFISH data processing and analysis

Spatial imaging data were processed using the MERSCOPE system and explored using the MERSCOPE visualizer software (v2.3, Qt version 5.15.2). Following initial processing steps on the MERSCOPE system, processed outputs were used to generate segmentation and expression count matrices using the Vizgen Postprocessing Tool, utilizing the watershed algorithm as implemented by scikit-image^43^ for segmentation, with seeds determined via the StarDist algorithm^44^. The configuration file for this process is made available at the link at the end of this paragraph. Notably, a set of patches had to be applied to VPT to resolve a deadlock within the tool stemming from the use of fork-based multiprocessing, resolved by utilizing spawn-based multiprocessing instead. This patch set is available at the link at the end of this paragraph. Following this initial image-based segmentation step, the resulting segmentations were further refined using Baysor version 0.6.2^45^, with a full configuration file available at the link at the end of this paragraph. Both previously described steps were run on Compute Canada infrastructure due to the very high computational requirements. Loom and geoJSON files generated by Baysor were used to generate Seurat objects containing expression and spatial data. Relevant cells were then annotated via spatial localisations using the open source tool scs (made available here: https://github.com/namemcguffin/scs). Following subsetting to only the relevant cells annotated previously, data was log-normalized, scaled, principal components were computed, UMAP embeddings were calculated, a nearest neighbour graph was generated, and clustering was conducted using the leiden algorithm via the API provided by the Seurat library^46^. Differential gene expression analysis was conducted also via Seurat using the FindMarkers function^46^. Visualisation for MERFISH data were generated by using the ggplot2 library directly. All software described above for this publication (including configuration files, patch sets, docker image definitions, scripts used to run pipelines on Compute Canada infrastructure, as well as processing and figure generation code) is available at the following link: https://github.com/stratton-lab/degenhard-2024-meninges_atlas.

