## Supplemental Figures for "An integrative layer-resolved atlas of the adult human meninges"

#### Supplemental Figure 1

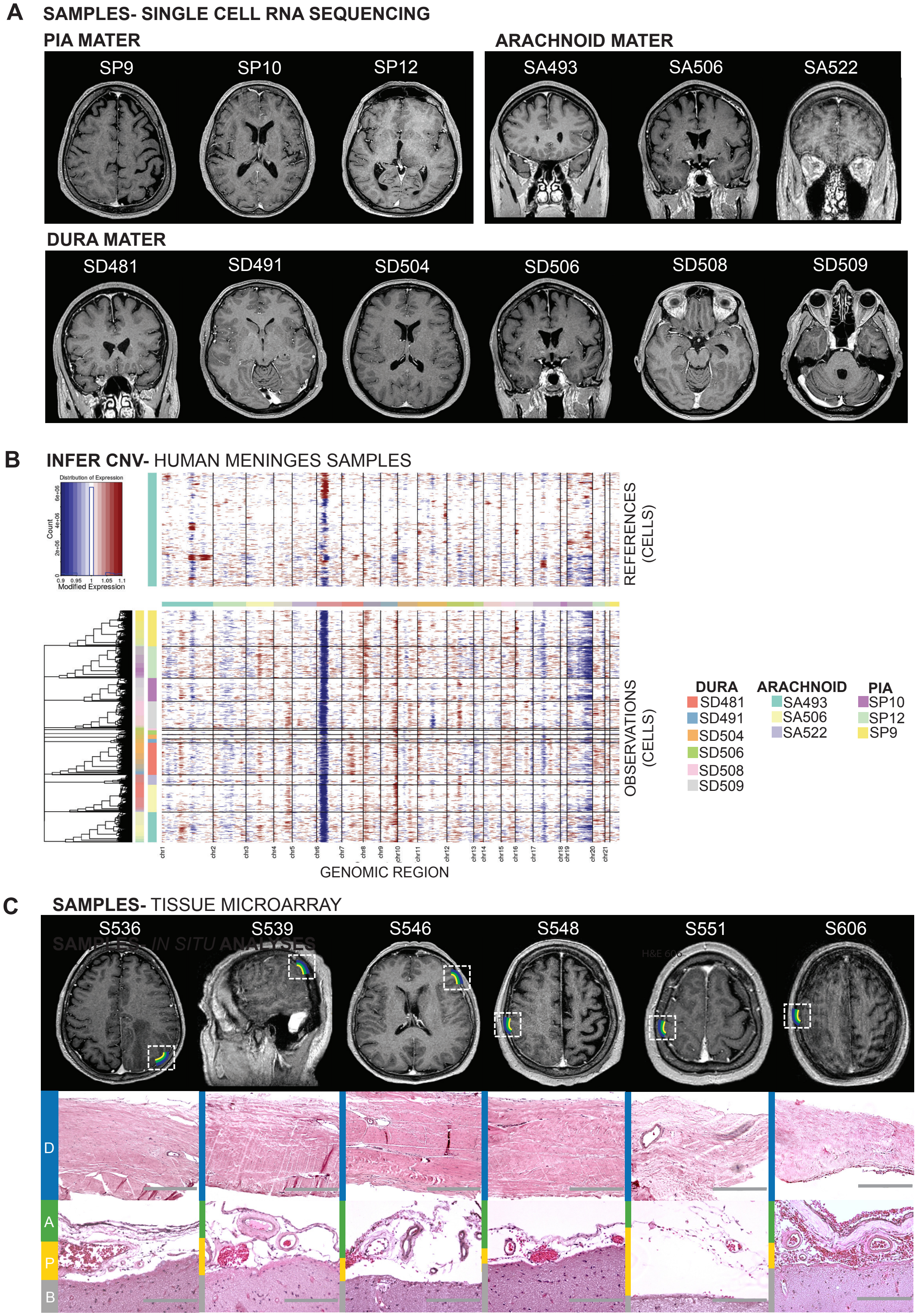

**Supplemental Figure 1: Patient-specific phenotyping with preoperative MRI, H&E staining, and CNV analysis confirms usage of healthy meninges tissue samples.**

(A) Preoperative MR images of patients undergoing resection of meninges used for scRNAseq. **(B)** Copy number variation analysis comparing immune cells (gene expression reference) to meningeal fibroblasts to detect any large scale copy number aberrations. Chromosome numbers are labelled. Red indicates gain of function and blue indicates loss of function. **(C)** Preoperative MR images of six patients used for tissue microarray preparation. Dashed white boxes indicate the location of meninges resection during surgery. H&E images demonstrate preservation of meningeal anatomy in the samples extracted from these patients. Scale bars = 400μm (dura); 200μm (leptomeninges).

### Supplemental Figure 2

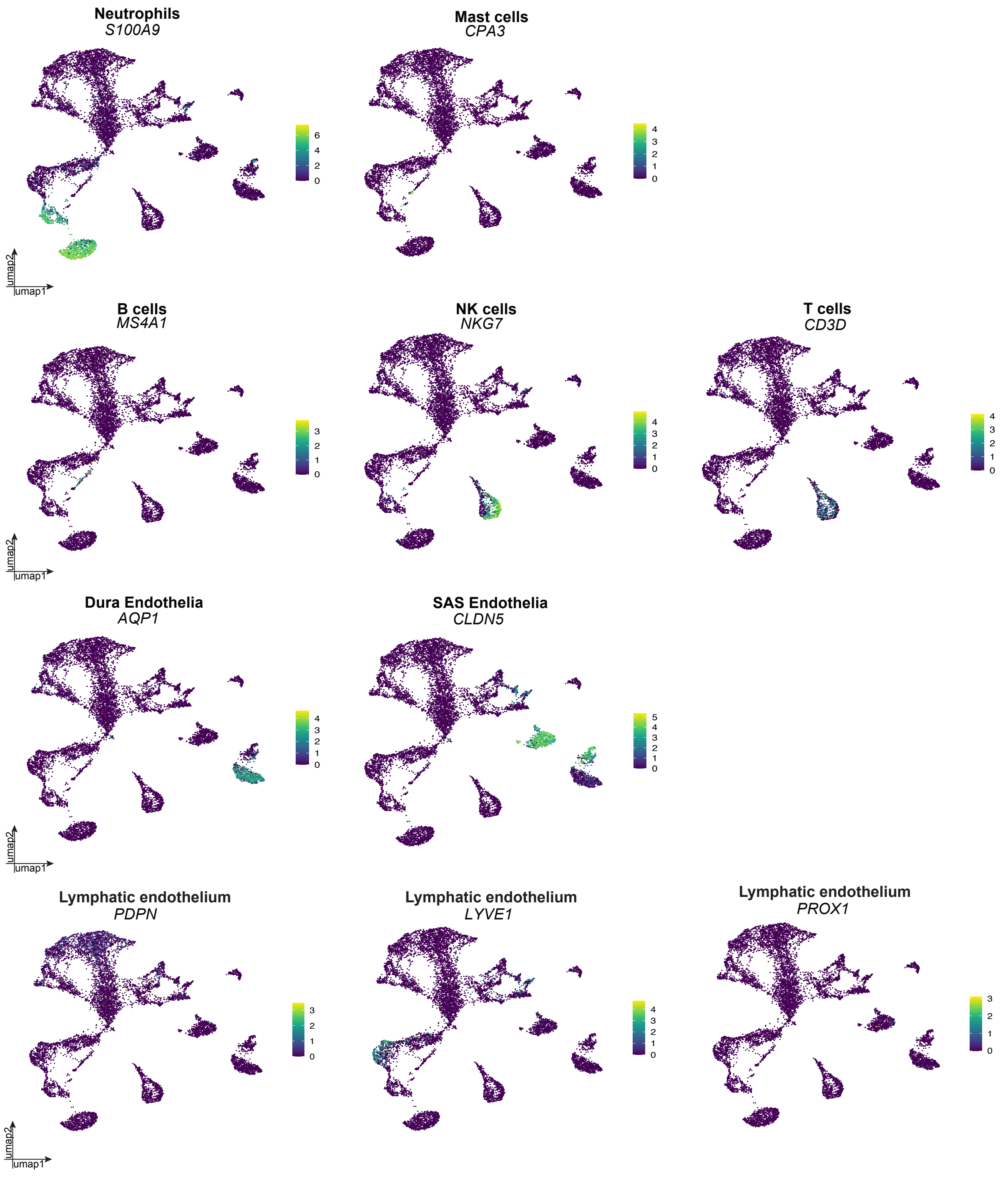

**Supplemental Figure 2: Feature plots of cell type-specific genes.**

Cell type-specific feature plots: T cells (*CD3D*), NK cells (*NKG7*), B cells (*MS4A1*), mast cells (*CPA3*), neutrophils (*S100A9*), SAS endothelia (*CLDN5*), dura endothelia (*AQP1*) and lymphatic endothelium (*PDPN, LYVE1, PROX1*).

#### Supplemental Figure 3

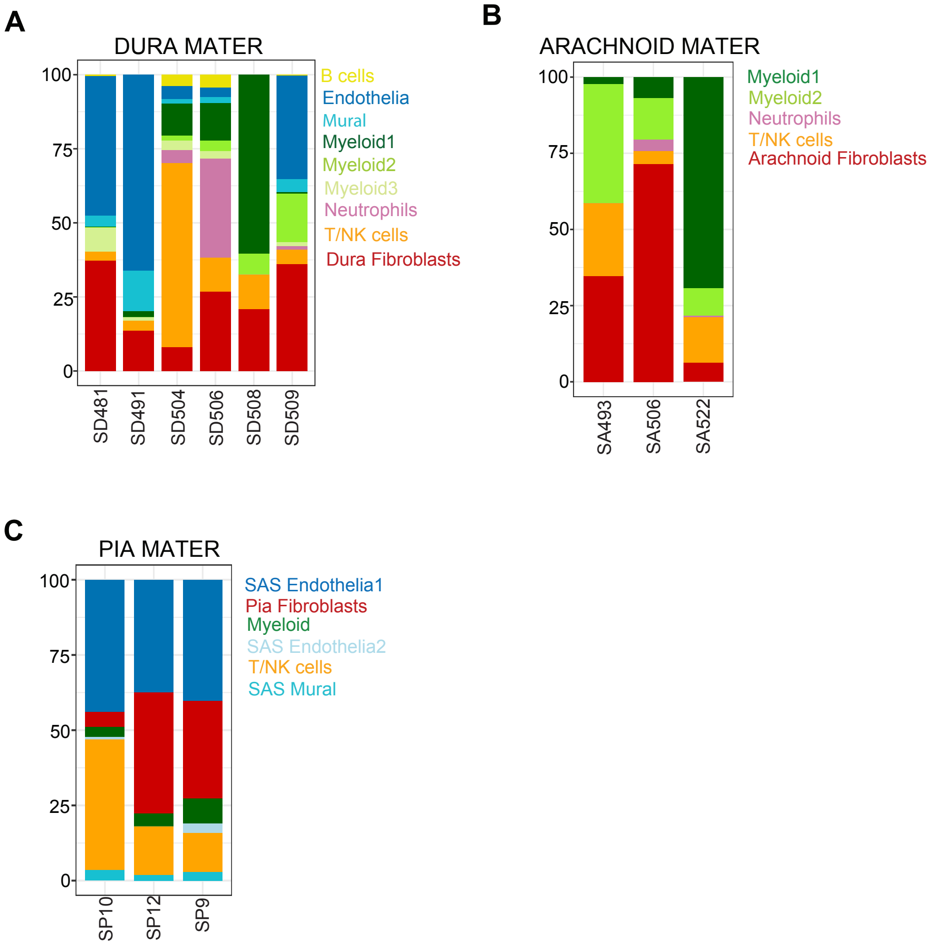

**Supplemental Figure 3: Layer-specific cell type proportions. (A)** Cell type proportions for six dura samples. **(B)** Cell type proportions for three arachnoid samples. **(C)** Cell type proportions for three pia samples.

#### Supplemental Figure 4

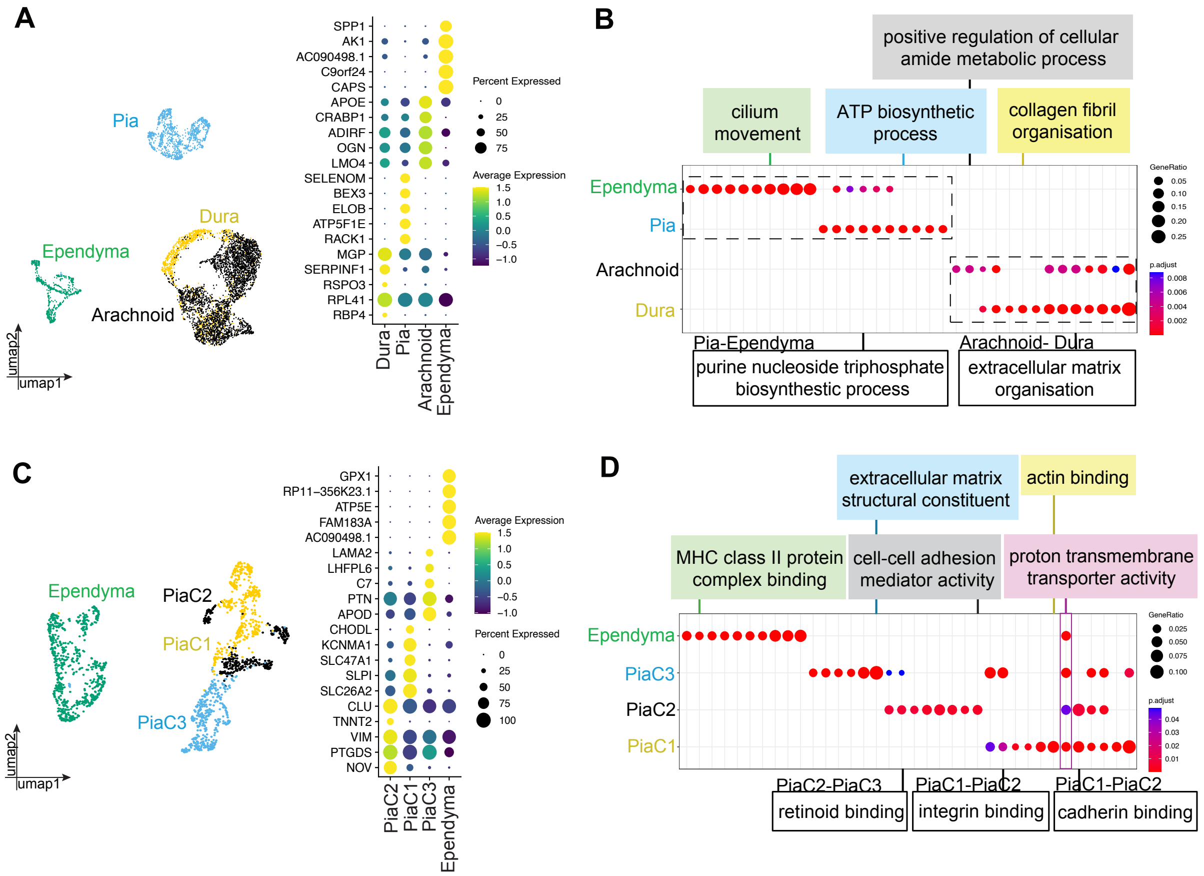

**Supplemental Figure 4: Integration of meningeal fibroblasts with ependymal cells reveals transcriptomic similarities between CSF-facing and brain-covering border cells. (A)** UMAP of layer-specific meningeal fibroblast populations (dura, arachnoid, and pia) and multiciliated ependymal cells. Adjacent dot plot demonstrating five key cell type marker genes for each cell population. Dot colour represents average expression level, and dot size represents the percentage of cells expressing the gene. **(B)** GO plot demonstrating representative biological pathways enriched in layer-specific meningeal fibroblasts and multiciliated ependymal cells compared to each other: dura fibroblasts (collagen fibril organization), arachnoid fibroblasts (cellular amide metabolic process), pia fibroblasts (ATP biosynthetic process), and ependymal cells (cilium movement). Select shared biological pathways are also shown: extracellular matrix organization (arachnoid fibroblasts-dura fibroblasts) and purine nucleoside triphosphate biosynthesis (pia fibroblasts-ependymal cells). Dot colour represents adjusted p-value, and dot size represents the gene ratio. **(C)** UMAP of pia fibroblast subpopulations (PiaC1, PiaC2, and PiaC3) and multiciliated ependymal cells with all ciliary genes removed. Adjacent dot plot demonstrating five key cell type marker genes for each cell population. Dot colour represents average expression level, and dot size represents the percentage of cells expressing the gene. (D) GO plot demonstrating representative biological pathways enriched in pia fibroblast subtypes and multiciliated ependymal cells (with cilia genes removed) compared to each other: PiaC1 fibroblasts (actin binding); PiaC2 fibroblasts (cell adhesion activity); PiaC3 fibroblasts (extracellular matrix structural constituent); and ependymal cells (MHC class II protein complex binding). Select shared biological pathways are also shown: cadherin binding (PiaC1-PiaC2); proton transmembrane transporter activity (PiaC1-PiaC2-PiaC3-ependymal cells); integrin binding (PiaC1-PiaC3); and retinoid binding (PiaC2-PiaC3).

#### Supplemental Figure 5

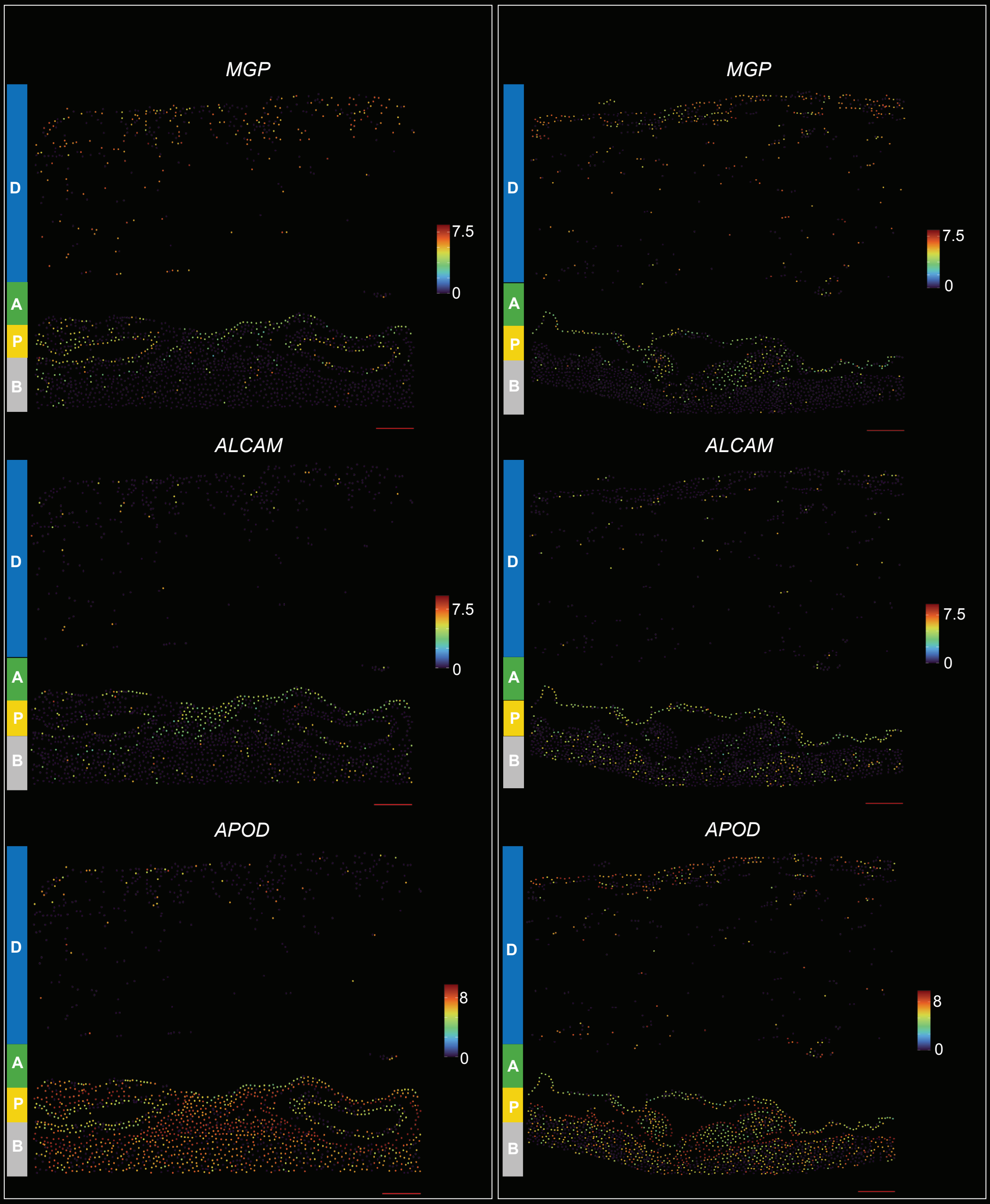


LEGEND

**D:** Dura
**A:** Arachnoid
**P:** Pia
**B:** Brain Parenchyma

**Supplemental Figure 5**: **MERFISH confirms scRNAseq-derived fibroblast-associated layer marker expression *in situ***. **(A)** MERFISH panels from two samples demonstrating elevated average *ALCAM* expression in arachnoid compared to the dura and pia. **(B)** MERFISH panels from two samples demonstrating elevated average *APOD* expression in the pia compared to the dura and arachnoid. **(C)** MERFISH panels from two samples demonstrating elevated average *MGP* expression in the dura compared to the arachnoid and pia. Dot colour indicates the average expression level of a particular gene. Scale bars = 100μm.

#### Supplemental Figure 6

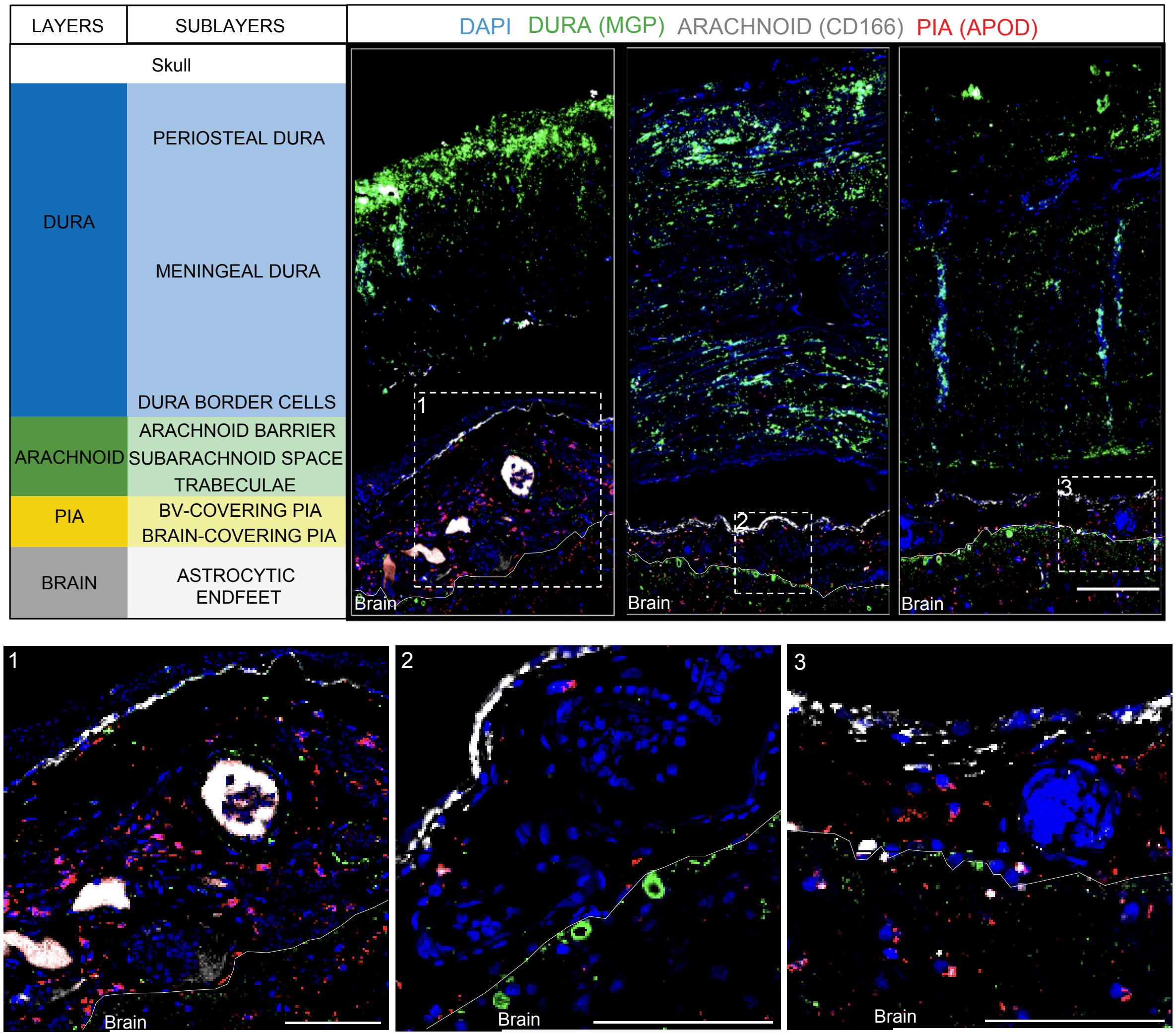

**Supplemental Figure 6**: **Protein Marker Labelling confirms scRNAseq-derived fibroblast-associated layer marker expression *in situ*.** (A) Protein immunolabelling of three fibroblast-associated meningeal layer markers: MGP (dura fibroblasts); CD166 (arachnoid fibroblasts); and APOD (pia fibroblasts). Meningeal architecture is annotated on the left hand side. Panels 1-3 represent three separate samples. Scale bar = 100μm. (B) Magnified images of the fibroblast-associated layer marker protein stains in panel A. Image locations are indicated by dashed white boxes in panel A. Scale bars = 100μm.

#### Supplemental Figure 7

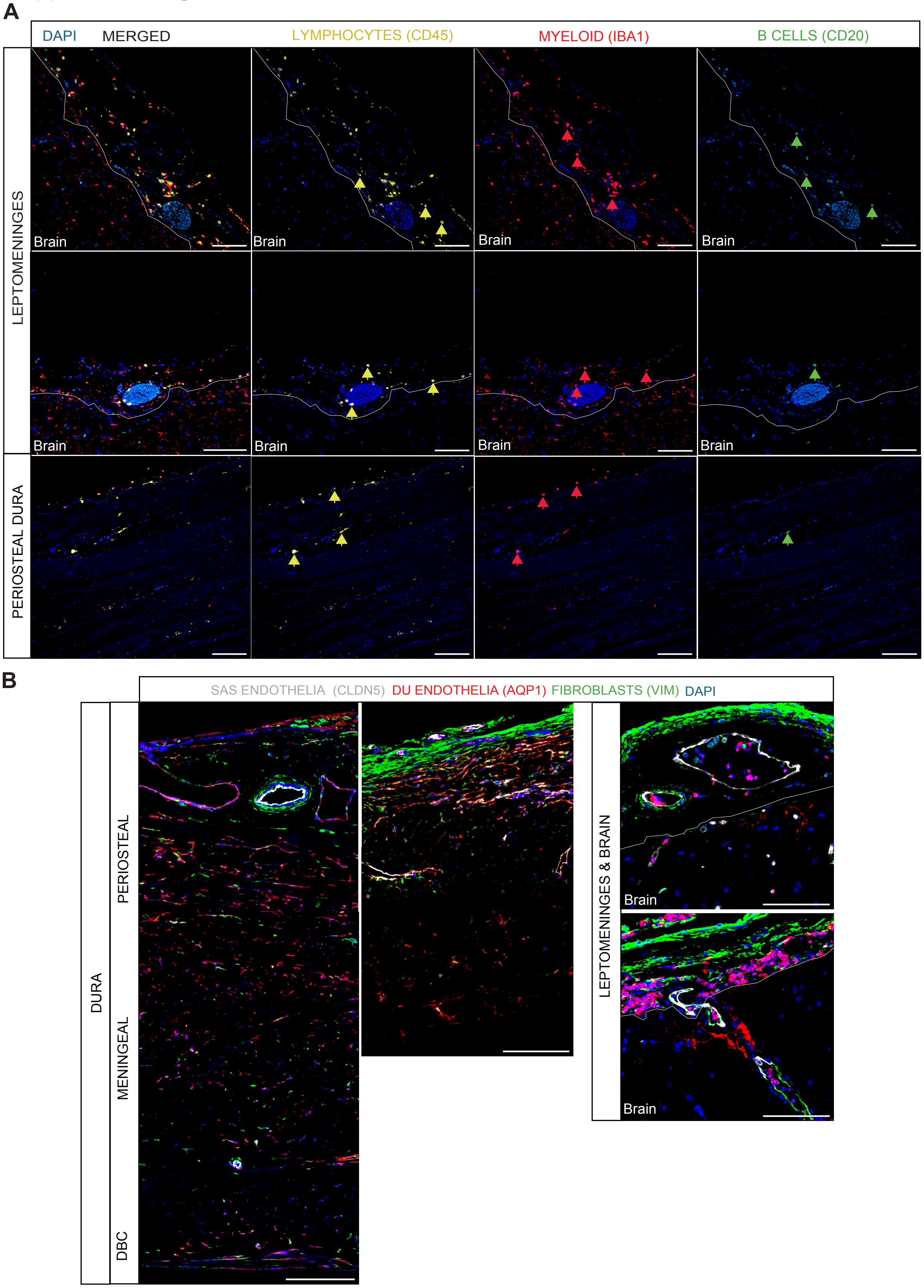

**Supplemental Figure 7**: **Protein Marker Labelling confirms the layer-specific spatial localisation of vascular and immune cell populations in the human meninges.** (A) Immune cell localization in the meninges: B cells (CD20); lymphocytes (CD45); and myeloid cells (Iba1). Arrows indicate examples of positively stained cells for respective markers. (B) Vascular marker expression of dura endothelia (AQP1) and SAS endothelia (CLDN5) with general fibroblast staining (VIM) demonstrates diverse vascular architecture in dura and leptomeninges. Scale bars = 100μm.

#### Supplemental Figure 8

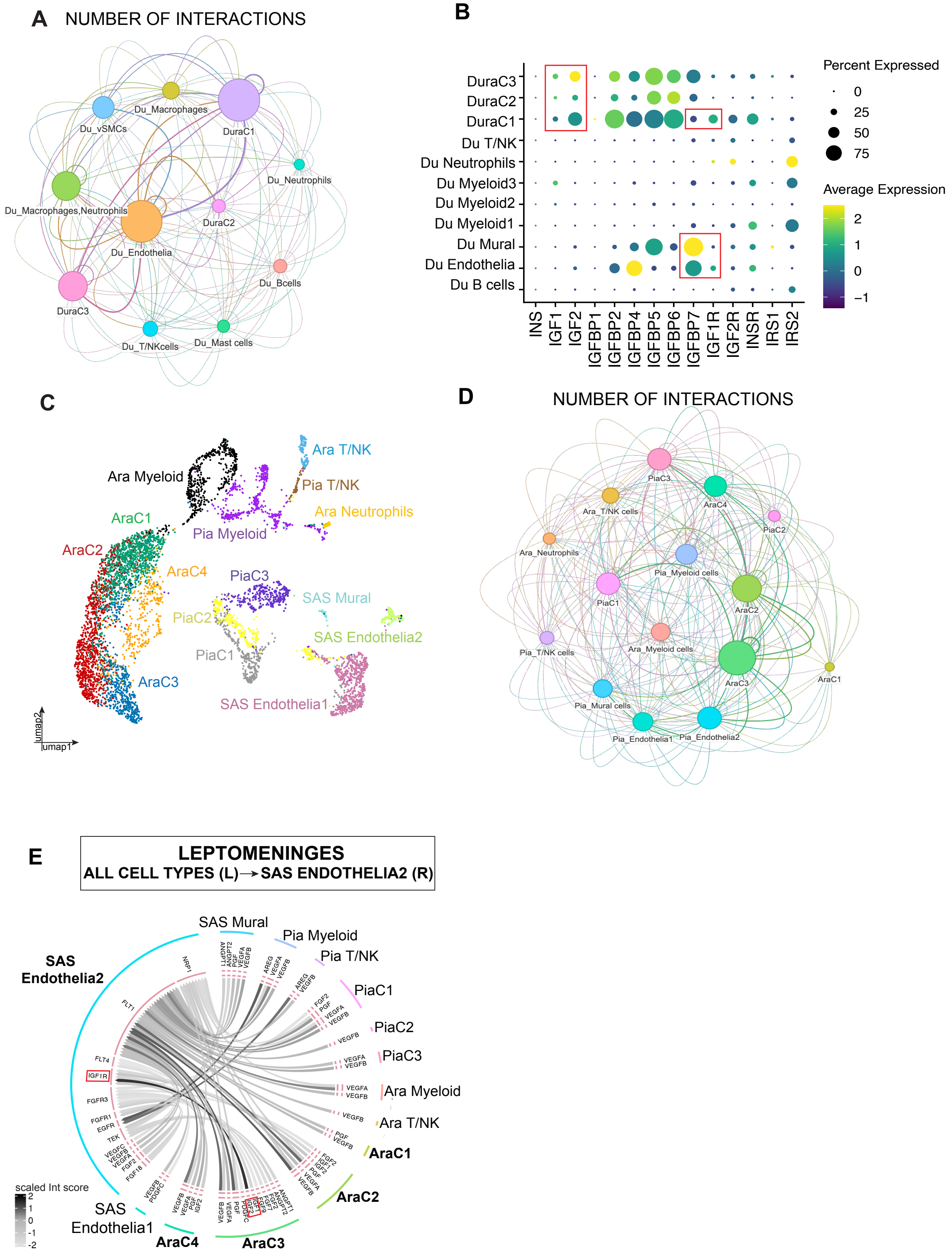

**Supplemental Figure 8**: **Cell-cell interaction analysis predicts VEGF- and IGF-mediated crosstalk between fibroblasts and endothelial cells of the dura and SAS**. **(A)** Interaction network plot of all cell types in the dura. Line colour is associated with cell type, and line thickness is indicative of the number of interactions. **(B)** Dot plot of canonical insulin growth factor signaling gene expression in dura cell types. Dot colour indicates average expression and dot size indicates percentage of cells expressing the gene. **(C)** Merged object of all cell clusters in the leptomeninges. **(D)** Interaction network plot of all cell types in the leptomeninges. Line colour is associated with cell type, and line thickness is indicative of the number of interactions. **(E)** Circle plot demonstrating the top significantly enriched ligand-receptor interactions pairs between all cell types in the leptomeninges and SAS Endothelia2. Line colour indicates scaled interaction score. **(F)** Circle plot demonstrating the top significantly enriched ligand-receptor interactions pairs between PiaC2 and cell types in the leptomeninges. Line colour indicates scaled interaction score.
